# Individual differences reveal distinct age and pubertal contributions to the refinement of the functional cortical hierarchy during adolescence

**DOI:** 10.64898/2026.05.07.723547

**Authors:** Bianca Serio, Lars Dinkelbach, Mylla Marsiglia, Laura Waite, Felix Hoffstaedter, Daniel S Margulies, Simon B Eickhoff, Sofie L Valk

**Affiliations:** Max Planck Institute for Human Cognitive and Brain Sciences, Leipzig, Germany; Max Planck School of Cognition, Leipzig, Germany; Institute of Neuroscience and Medicine, Brain & Behavior (INM-7), Research Centre Jülich, Jülich, Germany; Institute of Systems Neuroscience, Medical Faculty, Heinrich Heine University Düsseldorf, Düsseldorf, Germany; Department of Pediatrics II, University Hospital Essen, University of Duisburg-Essen, Essen, Germany; Institute of Sex- and Gender-sensitive Medicine, University Hospital Essen, University of Duisburg-Essen, Essen, Germany; Faculty of Life Sciences, Leipzig University, Leipzig, Germany; International Max Planck Research School on Cognitive Neuroimaging, Leipzig, Germany; Centre National de la Recherche Scientifique (CNRS), UAR 1329, Oxford, UK; Oxford Centre for Integrative Neuroimaging (OxCIN), FMRIB, Nuffield Department of Clinical Neurosciences, University of Oxford, Oxford, United Kingdom

## Abstract

The development of the functional cortical hierarchy, spanning sensorimotor to association systems, is exclusively studied as a function of age. During adolescence, this overlooks puberty as a major neurodevelopmental driver and source of variability. We studied sensorimotor-association axis refinement longitudinally (6323 observations across 4919 subjects), leveraging individual differences to disentangle chronological age from pubertal effects. We derived low dimensional features of sensorimotor-association axis development from resting-state functional connectomes, revealing substantial inter-individual heterogeneity in maturational trajectories that challenge group-level developmental trends and milestones. Then, we demonstrate independent effects of age and pubertal stage on sensorimotor-association axis refinement through the polarization of the cortical hierarchy. We further show that coordinated system-level shifts in network topology reflect an ongoing specialization of functional connectivity profiles across all major functional networks. Our findings frame adolescent hierarchical functional cortical maturation as an individualized, multifactorial phenomenon shaped by distinct chronological age and pubertal processes.

## Introduction

Adolescence is characterized by significant biological and psychosocial changes, representing a sensitive period for the onset of psychiatric disorders^1^ and for the development of functional brain organization subserving higher-order cognition^2,3^. It marks the transition from childhood to adulthood through puberty, a process of sexual maturation triggered by surges in the production of steroid hormones responsible for sex-specific physical and physiological bodily transformations^4^. Although pubertal development is loosely indexed by chronological age, these two processes represent different developmental mechanisms. Steroid hormones are important neuromodulators, and their pubertal increases act on structural and functional brain development through organizational and activational effects^5–9^. In turn, the neuro-developmental effects associated with chronological age globally reflect changes shaped by an interplay of (epi)genetic programming^10,11^ and experience-dependent plasticity^12–14^. Beyond the substantial collinearity between chronological age and pubertal stage, there is also significant variability in the timing of pubertal maturation between individuals and across sexes^15^, indicating that the two are not perfectly coupled – a dissociation that can be leveraged to disentangle their respective effects on brain development. Despite compelling evidence of age-independent pubertal effects on human brain structural maturation^16–23^ and emerging findings of pubertal associations with functional network connectivity^24,25^, the development of macroscale functional cortical organization has systematically been studied as a function of chronological age and largely focuses on group averages. Leveraging individual differences to disentangle the effects of chronological age and pubertal processes is therefore a crucial step towards understanding the mechanisms driving adolescent functional neurodevelopment.

A useful framework to study the development of functional brain organization is the sensorimotor-association (S-A) axis, which hierarchically differentiates unimodal sensorimotor systems, supporting primary visual and somatomotor functions, from transmodal association systems, supporting more abstract cognition. The S-A axis represents a major topographical principle of macroscale cortical organization, as it reflects a correspondence of variation in cortical features across modalities beyond functional connectivity, including also microstructure, gene expression, and receptor architecture, as well as developmental and evolutionary expansion^26–32^. On the one hand, the S-A axis has been described as a *spatiotemporal axis* of cortical development, characterizing the non-uniform and heterochronous maturation of cortical regions^13,31^. For instance, features of functional activity, including functional connectivity strength^33,34^ and intrinsic cortical activity^17,35^, display age-dependent patterns of maturation along this axis, with sensorimotor regions also developing earlier in childhood and association regions later during adolescence^13,31^. On the other hand, the S-A axis can be used as low dimensional *spatial embedding*, commonly known as a cortical gradient of brain organization, representing the primary axis of variation in resting-state functional connectivity in adults^29^. Along this gradient, cortical regions are organized in a spatially continuous order, whereby regions situated close to one another present similarities in their functional connectivity. Validated as a reliable and methodologically robust measure to characterize and study macroscale functional cortical organization^36^, the S-A axis has also been shown to capture both sex differences^37^ and effects of transient changes in steroid hormone levels^38^ in adults, suggesting its sensitivity to inter- and intra-individual sources of variability that are relevant to pubertal development. As such, the functional S-A axis as a spatial embedding represents a meaningful and unifying framework to study pubertal effects on a key principle of macroscale hierarchical cortical organization. Yet, its refinement during adolescence remains elusive, with its biological underpinnings unexamined, particularly the role of pubertal maturation.

Previous work has described group-level average changes in the development of the functional S-A axis as a function of age, reporting both a gradual refinement of sensorimotor-to-association spatial patterns from childhood to early adulthood and a discrete developmental reorganization of functional gradients^39,40^. Namely, the S-A axis –which in children is the secondary gradient of functional connectivity– has been shown to become the principal gradient explaining the most variance in connectivity patterns during adolescence. At the group level, this “gradient flip” is reported at around ages 11-12^39^ and 14-16^40^, depending on the study, and marks a categorical transition from child- to adult-like macroscale gradient organization. To describe this developmental transition at the individual level, gradual age-dependent increases in the variance explained by the S-A axis have been reported^41^, as well as changes in the time spent in dynamic intrinsic functional propagation states^42^. However, to enable inter-individual comparability, such studies relied on the orthogonal alignment of individual gradients to a shared dimensional space via a reference template. This poses a methodological challenge as it fundamentally distorts individual variability^43^, both in the continuous differentiation of S-A axis loadings and in the discrete gradient-level order of variance explained, which is fixed to that of the reference used for alignment. These continuous and discrete features are indeed both central to characterizing the transition to adult-like organization, although the degree to which they are biologically differentiable is unclear. To date, S-A axis maturation during adolescence therefore remains most reliably described by group-level averages as they do not resort to alignment, although they collapse the inter-individual variability in pubertal timing needed to disentangle chronological age from pubertal influences on S-A axis development.

To understand how underlying functional network maturation shapes the transition to adult-like gradient organization, virtual lesioning approaches have been used to highlight the key role of the ventral attention^44^ and default-mode^40^ networks in S-A axis refinement during adolescence. However, whether a single network has the specificity to drive such macroscale developmental changes is debatable. A hallmark of adolescent neurodevelopment is in fact the broad and distributed reorganization of functional networks^45–48^, illustrated by coordinated developmental shifts in within and between functional network connectivity^49–55^. Overall, the transition from childhood to adulthood is represented by decreasing global connectivity, suggesting gradual segregation of functional network communities^34,56^. These trends are reflected in an increasing range of S-A axis loadings during adolescence, indicating increasing differences in functional connectivity patterns between the two poles of the axis^40,41,57^. In parallel, from late childhood to early adulthood, changes in the S-A axis loadings of some functional networks shift them toward more extreme positions on the S-A axis^40,57^. However, such findings again relied on orthogonal alignment of individual gradients to a common template, introducing systematic biases by conflating individual developmental trajectories with distortions imposed by alignment and further disregarding the discrete transition to adult-like organization. The individual-level refinement of the S-A axis during adolescence thus remains an open question from both biological and methodological perspectives. The degree to which individual differences in functional network development reflect independent pubertal processes –separable from effects of chronological age– is also unclear, as is their significance in terms of changes in intrinsic functional connectivity. Addressing these gaps requires examining shifts of functional networks along the S-A axis from a systems perspective and at the individual level –free from template alignment constraints– while disentangling the contributions of age and puberty, and characterizing their intrinsic functional correlates. Doing so would yield a mechanistically grounded and methodologically rigorous explanatory account of hierarchical functional maturation during adolescence.

Here, we investigated the independent effects of chronological age and pubertal stage on the refinement of the functional cortical hierarchy during adolescence. To do so, we leveraged a system-level approach and focused on inter-individual differences in a large longitudinal cohort of 9- to 16-year-olds with up to three resting-state functional MRI scans each. Using alignment-free measures capturing both continuous and discrete low-dimensional features of S-A axis maturation, we reveal that the development of functional cortical organization is far more heterogeneous across individuals than group-level studies suggest, highlighting that group averages not only mask the variability of individual trajectories but may actively misrepresent the directions of developmental trends as well as the normative timing of developmental milestones for cortical reorganization. We demonstrate that chronological age and pubertal stage make distinct, additive contributions to the individual-level developmental trajectories of different features of hierarchical functional organization, operating through coordinated shifts in functional network topology that drive S-A axis expansion. We show that this polarization of the functional hierarchy, in turn, reflects an ongoing specialization of functional connectivity profiles across all major functional networks, underscoring that meaningful functional refinement during adolescence is not confined to association systems. Together, our findings frame adolescent functional brain maturation as a multifactorial phenomenon, shaped by both chronological age and pubertal processes, and demonstrate that capturing this distinction requires preserving the individual differences that define it.

## Results

To investigate how the functional cortical hierarchy refines across adolescence, we leveraged data from the Adolescent Brain Cognitive Development (ABCD) study^58^, a large cohort of youths aged 9 to 16 years (*N*_observations_ = 6323, including 4919 unique subjects), using a longitudinal design tracking resting-state functional cortical maturation across three timepoints: baseline (*n* = 4064), 2-year (*n* = 1296) and 4-year (*n* = 963) follow-ups. First, we computed novel alignment-free individual-level features of functional S-A axis development to quantify characteristic continuous and discrete maturational changes in macroscale gradient organization, comparing them to previous developmental patterns reported at the group-level. Our primary aim was then to tease apart the contributions of chronological age and pubertal stage to the developmental trajectories of these features of S-A maturation. To do this, we relied on individual differences in pubertal timing by including both chronological age and pubertal stage as covariates in our models to isolate their relative contributions. Furthermore, to provide an explanation for S-A axis refinement at the system level, we examined whether topological shifts of functional networks underpin S-A axis expansion. Finally, we probed the functional underpinnings of S-A axis polarization in terms of changes in intrinsic functional connectivity profiles, leveraging the lengths of functional connections as measured by geodesic distances on the cortical surface.

We used Big Additive Models (BAMs) to capture non-linear trajectories of functional S-A axis development, which are a Generalized Additive Model (GAM) approach optimized for large-scale datasets. These models systematically accounted for the effects of major factors associated with brain development, including chronological age, pubertal stage, sex, and total surface area as the measure of brain size most strongly associated with variability in the functional S-A axis^37^. Models also included random nested effects, accounting for repeated measures, family membership (siblings) and research site, in order to control for the nested structure of the ABCD dataset. Given that pubertal timing is largely sex-dependent, with females undergoing puberty 1-1.5 years earlier than males on average^15^, we systematically tested for sex effects and visualized maturational trajectories by sex, in order to probe their sex-specificity. Finally, we conducted supplementary analyses, which included assessing the relationship between our individual-level measures of S-A axis maturation (Supplementary Fig. 1) and exploring effects related to changes in functional connectivity strength (Supplementary Fig. 2). To ascertain the specificity and robustness of our findings, we also performed sensitivity analyses including alternative methods for characterizing S-A expansion (Supplementary Fig. 3), the inclusion of potential confounders in our statistical models, such as socioeconomic status^59^ (SES; specifically, parental education; Supplementary Fig. 4) and body mass index^60^ (BMI, z-scored for age and sex; Supplementary Fig. 5), and an alternative method for defining categories of lengths of functional connections (Supplementary Fig. 6), all of which confirmed the robustness of our results.

### Individual-level gradients reveal heterogeneity that challenges group-level accounts of S-A axis development

Studies commonly report descriptive, group-level averages of S-A axis measurements. These are computed by using eigendecomposition to derive typically ten low dimensional axes of variation in mean functional connectivity patterns (averaged across subjects), commonly known as cortical gradients of brain organization (G1-G10), ordered by their explained variance. Previous findings consistently report that the S-A axis is represented by the primary gradient (G1) explaining the most variance in adults^29,39,40,44^, whereas it is represented by the secondary gradient (G2), explaining the second most variance in children^39,40,44^.

To anchor our study in previous findings, we first computed group-level cortical gradients of brain organization from average functional connectivity matrices on the Schaefer 400 parcellation^61^ at the three available timepoints of data collection. Contrary to reports of gradual group-level refinement during childhood and adolescence^39,40,44^, we found that the spatial patterns of loadings in mean G1 and G2 remained largely invariant across the 9- to 16-year age range, as established with spin-based spatial permutation testing^62^ of pairwise correlations across timepoints (G1, *r* = 0.979-0.995, *p*_spin_ < .001; G2, *r* = 0.988-0.996, *p*_spin_ < .001; **Fig. 1A,B**). The relative variance explained by these top two gradients also remained stable over time (**Fig. 1C**). Notably, at the mean group level, the S-A axis was consistently represented by the second gradient (G2), rather than shifting to the primary gradient (G1) – a discrete transition to adult-like patterns of organization that we will refer to as the ‘gradient flip’. This lack of group-level gradient flip across our 9- to 16-year window –suggesting its occurrence after the latest measured timepoint– deviates from previous evidence reporting its occurrence between ages 11 and 12^39^ or 14 and 16^40^, depending on the study.

**Figure 1.**
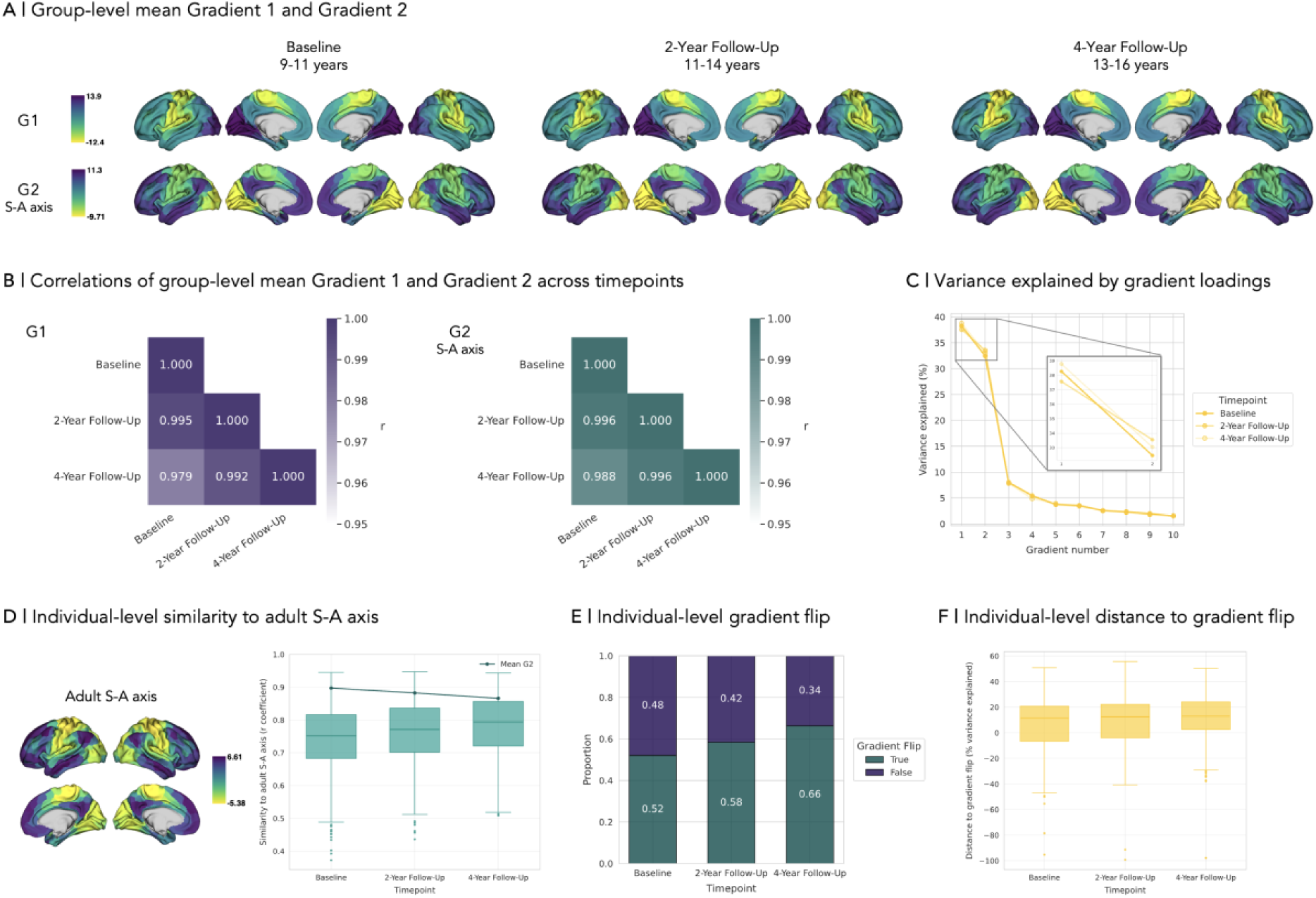
Differences in features of sensorimotor-association (S-A) axis development when computed at the group- and individual-level. **A |** Group-level mean Gradient 1 (G1) and Gradient 2 (G2) of functional cortical organization by timepoint, where G2 reflects S-A axis patterns across timepoints; **B |** Spearman’s correlation coefficients (absolute values) of group-level mean G1 and G2 loadings across timepoints, with statistical significance corrected for spatial autocorrelation using spin permutation testing^62^ (G1, *r* = 0.979-0.995, *p*_spin_ < .001; G2, *r* = 0.988-0.996, *p*_spin_ < .001); **C |** Variance explained by the loadings of group-level mean gradients 1-10 by timepoint, focusing on G1 and G2; **D |** Distribution by timepoint of individual-level similarity to the adult S-A axis, defined as the association (Spearman’s correlation coefficient) between subject-level S-A axis and an adult group-level mean S-A axis template (data from ref.^29^), ranging *r* = 0.37-0.95 across timepoints, with overlayed mean G2 line showing the correlations between group-level mean S-A axis by timepoint and the adult S-A axis template; **E |** Proportions by timepoint for individual-level gradient flip, capturing shifts in the relative variance explained by the S-A axis, i.e., whether S-A patterns are reflected in G1 (True) or not (False). **F |** Distribution by timepoint of individual-level distance to gradient flip, defined as the difference in scaled variance explained (%) between the S-A axis and the highest-variance non-S-A gradient, where negative values indicate that the gradient flip has not yet occurred and positive values indicate that the S-A axis is the gradient explaining the most variance (i.e., G1). Boxplots are represented by boxes extending from the first to the third quartiles of the data with a line representing the median, whiskers extending from the boxes to the farthest data points within 1.5 times the interquartile range, and fliers representing data points past the ends of the whiskers.

We next computed cortical gradients for each subject –following the eigendecomposition procedure outlined above– in order to capture individual differences in S-A axis development. Crucially, to preserve variability across subjects, we did not align individual-level gradients to any group average. By identifying the gradient that most strongly correlated with an adult group-level mean S-A axis template^29^, we quantified two novel key features of S-A axis development for each subject: (1) the similarity to the adult S-A axis, a continuous measure quantified by the Spearman correlation coefficient of the individual’s S-A axis loadings to the adult S-A axis template, and (2) the occurrence of the gradient flip, a discrete, binary measure quantifying whether or not S-A patterns were reflected in G1 for that given individual, representative of the transition toward adult-like organization.

At the individual level, we observed substantial variability that is masked in group-level averages, as well as diverging trends and representations of milestones between individual- and group-level summaries. While group-level similarity to the adult template surprisingly slightly decreased across timepoints (mean G2; *r*_baseline_ = 0.90 to *r*_4y_ = 0.87), individual-level similarity showed a global upward trend (median S-A axis; *r*_baseline_ = 0.75 to *r*_4y_ = 0.79; **Fig. 1D**). The opposing directions of these trends underscore a fundamental divergence between group- and individual-level descriptions of continuous S-A axis maturation. Furthermore, the individual-level computation of the discrete gradient flip revealed that S-A axis patterns were already reflected in G1 in more than half of the sample at baseline (52%) and continued to increase across timepoints (58% at 2-year follow-up, 66% at 4-year follow-up; **Fig. 1E**). These results indicate that the discrete shift toward adult-like gradient organization is already underway in early adolescence in the majority of subjects in our sample – a key milestone that is obscured when data are averaged at the group level. To verify that the gradient flip occurrence does not merely reflect trivial differences in variance explained, we quantified a “distance to gradient flip” metric, defined as the difference in scaled variance explained between the S-A axis and the highest-variance non-S-A gradient, with negative values indicating that the gradient flip has not yet occurred and positive values indicating that the S-A axis is the gradient explaining the most variance. This metric revealed substantial inter-individual variability and a progressive increase in S-A axis dominance across timepoints (**Fig. 1F**), confirming that the discrete gradient flip is a milestone that occurs on the back of a meaningful and progressive developmental shift rather than an arbitrary relative difference in variance explained. Finally, we explored the interdependence between our features of S-A axis development, revealing a statistically significant relationship between individual similarity to the adult template and the probability of observing the gradient flip (Supplementary Fig. 1).

Collectively, these findings reveal substantial individual differences in the refinement of the S-A axis during adolescence, further underscoring that individual- and group-level summaries may paint a fundamentally different –at times opposing– picture of S-A axis maturation. For instance, group-level averages display trends of decreasing similarity to the adult S-A axis contrary to an increasing trend shown at the individual level. Furthermore, individuals seem to present adult-like hierarchical organization earlier than group-level analyses suggest, with group averages failing to reflect that the gradient flip already occurred in a majority of participants at baseline, a proportion that progressively increases across timepoints. This indicates that averaging across individuals not only obscures the heterogeneity of individual developmental trajectories but may misrepresent the normative trends and timing of cortical reorganization. These findings underscore the necessity of individual-level approaches to accurately capture the pace and individual variability of adolescent S-A axis maturation.

### Chronological age and pubertal stage show distinct additive effects on S-A axis development

Previous studies have characterized the refinement of the S-A axis during adolescence as a function of chronological age. However, given the key role of puberty in adolescent development, and considering that pubertal timing differs between individuals and across sexes^15^, we aimed to test the independent contributions of chronological age and pubertal stage using a sex-sensitive approach. Within our cohort, males were marginally older than females across all timepoints (**Fig. 2A**; *p* < 0.05 at baseline and 2-year follow-up). Conversely, females exhibited a more advanced pubertal stage based on physical pubertal characteristics, as measured by the Pubertal Development Scale (PDS; **Fig. 2B**; *p* < .001 at all timepoints)^63^. While chronological age (in months) and pubertal stage (PDS score) were moderately correlated (Spearman’s *r* = 0.59, *p* < .001; **Fig. 2C**), the Variance Inflation Factor (VIF = 1.96) indicated sufficient independent variance of age and pubertal stage beyond their collinearity for models to reliably estimate the unique effects of both covariates while statistically controlling for the other.

**Figure 2.**
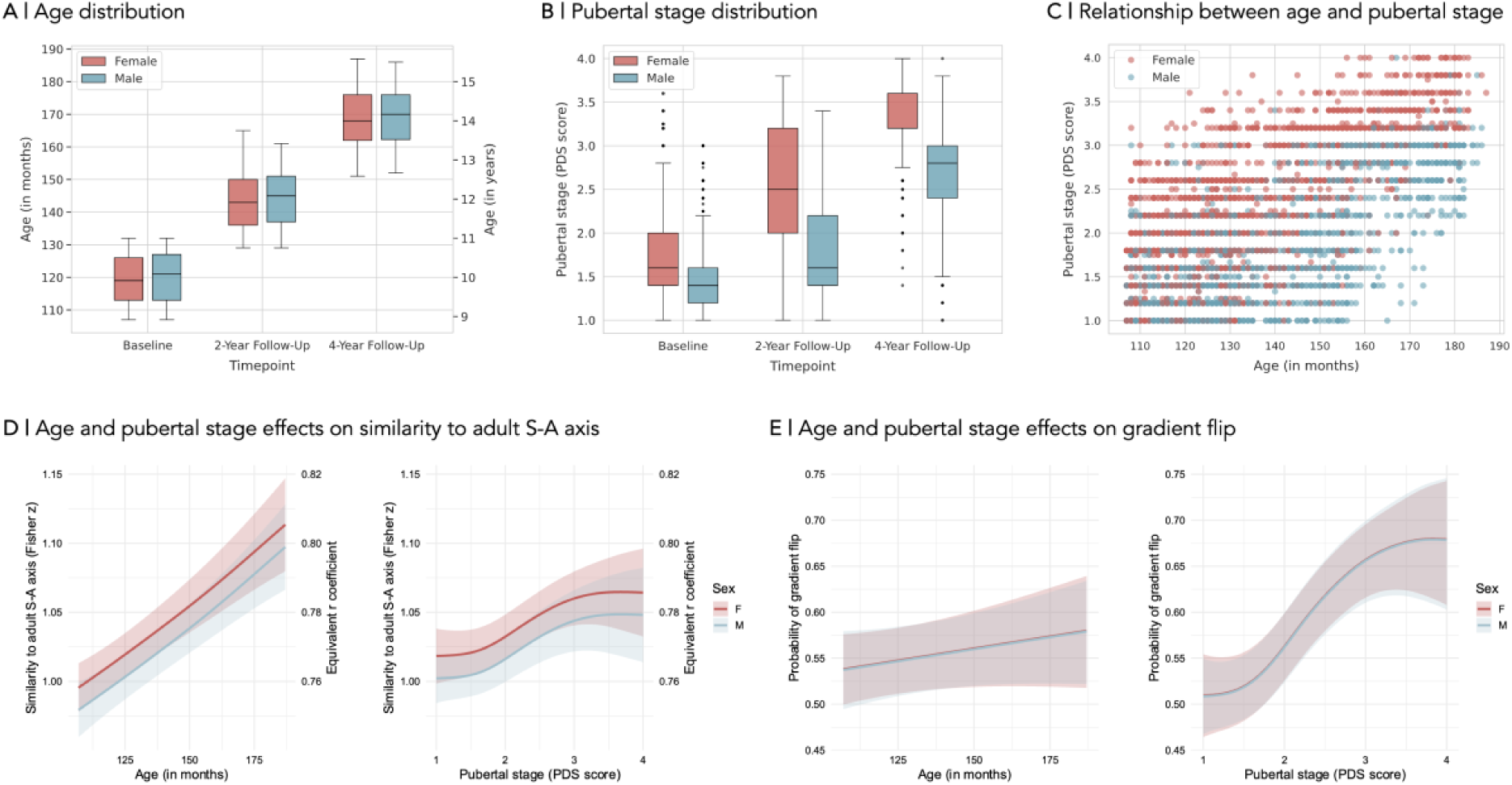
Distinct chronological age and pubertal stage effects on features of sensorimotor-association (S-A) axis development. **A |** Distribution of age (in months and years) by sex and timepoint; **B |** Distribution of pubertal stage score by sex and timepoint, characterizing physical pubertal development based on the Pubertal Development Scale (PDS) parent report, with scores ranging from 1 (indicating that pubertal development has not yet begun) to 4 (indicating that pubertal development is completed); **C |** Scatterplot displaying the relationship between age and pubertal stage, tested by Spearman correlation, *r* = 0.59, *p* < .001, color coded by sex; **D |** Predicted marginal trajectories for age and pubertal stage effects on the similarity to adult S-A axis by sex (model estimates are shown on the Fisher *r*-to-*z* transformed scale; a secondary y-axis displays back-transformed Spearman *r* values to ease interpretation); **E |** Predicted marginal trajectories for the age and pubertal stage effects on the probability of observing the gradient flip by sex (age effect is not statistically significant). Boxplots are represented by boxes extending from the first to the third quartiles of the data with a line representing the median, whiskers extending from the boxes to the farthest data points within 1.5 times the interquartile range, and fliers representing data points past the ends of the whiskers. Statistical effects were computed with Big Additive Models (BAMs) including fixed effects of chronological age, pubertal stage, sex and total surface area, and random nested effects (subject, family, site). Predictions were generated by holding all continuous covariates at their mean values and by excluding random nested effects to produce population-level trajectories. Shaded ribbons represent 95% Bayesian credible intervals derived from the posterior distribution of the model coefficients, representing the uncertainty in the population-level response by excluding the variance attributed to the random nested effects of subject, family, and site.

We first tested for the effects of chronological age and pubertal stage on the similarity to the adult S-A axis. Both variables uniquely contributed to increases in S-A axis maturation (full model deviance explained = 38.5%) and their effects were represented by distinct trajectories. The effect of age was near-linear and robust (*EDF* = 1.37, *F* = 11.44, *p* < .001; **Fig. 2D, left**), whereas pubertal stage followed a non-linear and flatter sigmoidal trajectory (*EDF* = 2.16, *F* = 2.11, *p* = 0.002; **Fig. 2D, right**). This sigmoidal effect was characterized by plateaus at early and late puberty (PDS < 1.5 and > 3.0), with an accelerated increase in adult-like similarity specifically during mid-puberty. While females showed a constant, slightly greater overall similarity to the adult S-A axis than males (*t* = -2.25, *p* = 0.025), we found no significant age-by-sex nor pubertal stage-by-sex interactions, highlighting a shared developmental trajectory across sexes.

In contrast, the probability of observing the gradient flip was robustly driven by pubertal stage (*EDF* = 2.53, χ^2^ = 27.73, *p* < .001; **Fig. 2E, right**) but not by age (*EDF* = 0.63, χ^2^ = 1.78, *p* = 0.11; **Fig. 2E, left**) (full model deviance explained = 14.2%). The effect of pubertal stage again followed a sigmoidal curve with a steep increase in gradient flip probability during the mid-pubertal window. Although there was no difference between the sexes in gradient flip probability (*z* = -0.05, *p* = 0.963) –and while the pubertal effect on the gradient flip was sex-independent, as evidenced by a lack of pubertal stage-by-sex interaction (*EDF* = 0.23, χ^2^ = 0.26, *p* = 0.189)– we did observe a significant age-by-sex interaction (*EDF* = 1.25, χ^2^ = 5.08, *p* = 0.014). Post-hoc analyses indicated that age marginally contributed to the gradient flip probability in males (*EDF* = 1.25, χ^2^ = 5.13, *p* = 0.013) but not in females (*EDF* = 0.00, χ^2^ = 0.00, *p* = 0.772), despite their substantially overlapping developmental trajectories.

Altogether, we show that chronological age and pubertal stage distinctly and additively contribute to different low dimensional features of functional hierarchical development across the sexes despite their temporal collinearity. These findings further highlight that the continuous and discrete features of S-A axis refinement –the similarity to adult S-A axis and the gradient flip– are biologically differentiable in providing complementary perspectives on S-A axis development. The pubertal transition appears to uniquely affect the categorical shift to adult-like gradient organization, where the S-A axis becomes the principal gradient explaining the most variance in functional connectivity. Altogether, mid-puberty presents as a window of enhanced cortical reorganization, a process that appears largely independent of sex, despite sex differences in pubertal timing.

### Coordinated topological network shifts drive S-A axis expansion and refinement

During adolescence, trends of global segregation of functional network communities have been previously reported^34,56^ and are reflected in an increasing range of S-A axis loadings^40,41,57^. To understand S-A axis maturation from a systems perspective, we investigated whether topological shifts in functional networks underpin S-A axis expansion while also assessing the independent contributions of chronological age and pubertal stage to this reorganizational process. Network dispersion in gradient space has been recently characterized using a centroid-based approach^37,38,64^, identifying the center of functional networks as their median loadings and quantifying distances between network centroids. Building on this approach, we derived a novel, individual-level measure of S-A axis expansion. This measure was obtained by computing the aggregate hierarchical distance of network centroids from the S-A axis origin, that is, by calculating the sum of squared median loadings of the seven Yeo networks^65^. This metric captures the degree to which functional communities diverge from the hierarchical center of the axis toward the sensorimotor and association poles, reflecting the global segregation of networks at the system level. S-A axis expansion outliers were excluded from our analyses, but sensitivity analyses confirmed that this expansion measure and its related effects are robust to methodological scale and network definition (i.e., when expansion is computed at the regional instead of the network level), and that results are independent of outlier exclusion (Supplementary Fig. 3).

We observed substantial heterogeneity across individuals in their network centroids, as well as a descriptive trend of increased polarization of functional networks across timepoints (**Fig. 3A**). Sensorimotor-leaning networks (visual and somatomotor networks), as well as transitional dorsal and ventral attention networks situated closer to the axis origin –all anchored in the negative half of the axis– displayed a shift toward more extreme negative loadings over time. Conversely, association-leaning networks anchored in the positive half of the axis (limbic, frontoparietal, and default-mode networks) shifted toward more extreme positive loadings. These divergent shifts represent an increasing segregation between sensorimotor and association networks and reflect a global increase in S-A axis expansion by timepoint, which also displayed high levels of variability across individuals (**Fig. 3B**).

**Figure 3.**
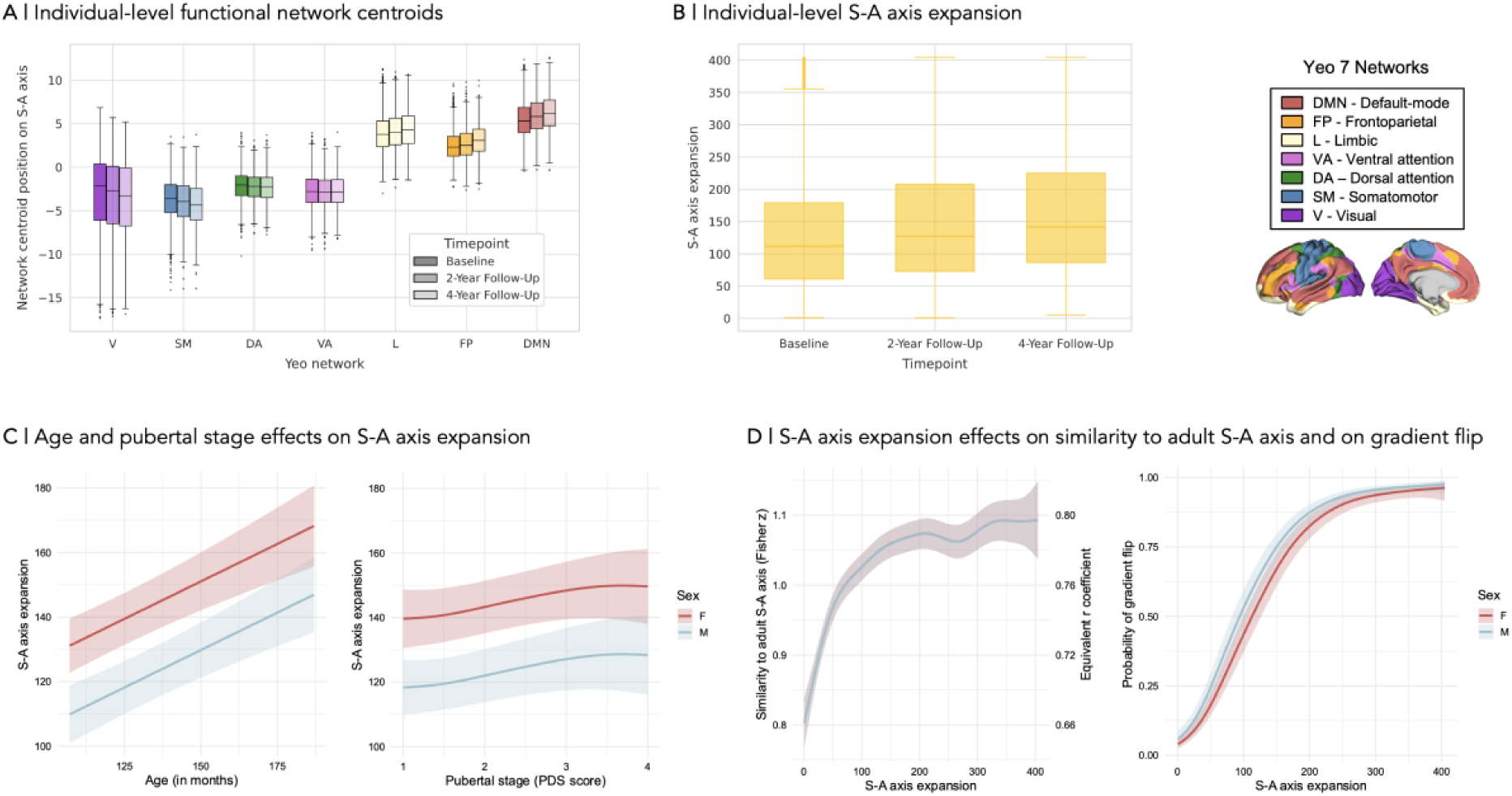
System-level topological shifts in functional networks driving sensorimotor-association (S-A) axis expansion and development. **A |** Distribution by timepoint of individual-level functional Yeo network centroids, representing the median of loadings across regions belonging to each of the seven Yeo networks; **B |** Distribution by timepoint of individual-level S-A axis expansion, defined as the sum of squared network centroids to capture total magnitude of dispersion from S-A axis origin (0); **C |** Predicted marginal trajectories for age and pubertal stage effects on S-A axis expansion by sex; **D |** Predicted marginal trajectories for the S-A axis expansion effect on the similarity to the adult S-A axis (model estimates are shown on the Fisher *r*-to-*z* transformed scale; a secondary y-axis displays back-transformed Spearman *r* values to ease interpretation) and on the probability of observing the gradient flip, by sex. Boxplots are represented by boxes extending from the first to the third quartiles of the data with a line representing the median, whiskers extending from the boxes to the farthest data points within 1.5 times the interquartile range, and fliers representing data points past the ends of the whiskers. Statistical effects were computed with Big Additive Models (BAMs) including fixed effects of chronological age, pubertal stage, sex and total surface area, and random nested effects (subject, family, site). Predictions were generated by holding all continuous covariates at their mean values and by excluding random nested effects to produce population-level trajectories. Shaded ribbons represent 95% Bayesian credible intervals derived from the posterior distribution of the model coefficients, representing the uncertainty in the population-level response by excluding the variance attributed to the random nested effects of subject, family, and site. V, visual; SM, somatomotor; DA, dorsal attention; VA, ventral attention; L, limbic; FP, frontoparietal; DMN, default-mode network.

We then probed the relative effects of chronological age and pubertal stage on S-A axis expansion, again finding distinct contributions of both factors (full model deviance explained = 48.7%). Age showed a robust, near-linear association with S-A axis expansion (*EDF* = 0.97, *F* = 9.86, *p* < .001; **Fig. 3C, left**) while pubertal stage followed a flatter sigmoidal trajectory (*EDF* = 0.85, *F* = 1.18, *p* = 0.019; **Fig. 3C, right**), with a subtle increase in the mid-pubertal window. Females exhibited slightly greater expansion overall (*t* = -7.29, *p* < .001), but developmental trajectories were consistent across sexes, as evidenced by no statistically significant age-by-sex (*EDF* = 0.00, *F* = 0.00, *p* = 0.919) nor pubertal stage-by-sex (*EDF* = 0.62, *F* = 0.22, *p* = 0.152) interaction effects on S-A axis expansion.

Critically, S-A axis expansion significantly contributed to the refinement of the functional hierarchy, predicting our two main features of S-A axis development above and beyond the previously reported age and pubertal stage effects (full model deviance explained = 36.8%). Expansion showed a biphasic effect on the similarity to the adult S-A axis (*EDF* = 6.70, *F* = 69.18, *p* < .001; **Fig. 3D, left**), with an initial phase of rapid maturation during early expansion followed by stabilization at high expansion values. Similarly, expansion significantly predicted the probability of the gradient flip (*EDF* = 4.57, χ^2^ = 1827.01, *p* < .001; **Fig. 3D, right**), following a sigmoidal curve characterized by a slight increase from 0.0 probability of gradient flip at very low levels of expansion, followed by a steep increase, and reaching a plateau near 1.0 at high expansion values. While there was no evidence of an expansion-by-sex interaction effect on the similarity to the adult S-A axis (*EDF* = 0.00, *F* = 0.00, *p* = 0.868), a marginal expansion-by-sex interaction in the gradient flip probability suggested a slightly steeper increase in males (*EDF* = 1.62, χ^2^ = 2.12, *p* = 0.037), though this interaction was not robust to sensitivity testing, namely when S-A axis expansion was computed at the regional level.

Together, these findings portray S-A axis expansion as a fundamental developmental characteristic of hierarchical functional organization that is sensitive to distinct effects of chronological age and pubertal stage. By quantifying the increasing system-level segregation between the sensorimotor and association poles, we found that sensorimotor-leaning networks become more sensorimotor-like and association-leaning networks more association-like during adolescence, thus refining the functional hierarchy. Furthermore, the observed non-linear biphasic effects of S-A axis expansion on the similarity to the adult S-A axis and gradient flip probability suggest a point of developmental saturation, highlighting that the macroscale functional hierarchy appears to stabilize into its adult-like configuration once a critical threshold of expansion is reached.

### The specialization of functional connectivity profiles across all major networks underpins S-A axis expansion

The above results show that S-A axis expansion –a phenomenon that was previously only described as increases in the range of S-A axis loadings^40,41,57^– is actually driven by hierarchical and system-level topological shifts of functional networks. Next, we sought to explain what developmental changes this expansion may represent at the intrinsic functional level. Previous work shows that low-dimensional S-A axis loadings predominantly reflect patterns of functional connectivity profiles, namely which regions are most strongly connected^37^. At the same time, a relevant feature of intrinsic functional connectivity is the characteristic length of connections that a region makes – more specifically the distance between a region and its strongest functionally connected partners on the cortical surface. Hypothesized to emerge from structural constraints^66^, these lengths of functional connections are highly related to the geometry of the cortex, as captured by geodesic distances along the cortical mantle, following gyrification^28,29,67^. Namely, geodesic distances between strongly functionally connected regions closely follow the S-A hierarchy, with sensorimotor and association systems predominantly making short- and long-range connections, respectively^67–69^. We therefore hypothesized that as the S-A axis expands –reflecting a global segregation of functional network communities in the maturing cortical hierarchy– networks would exhibit increasingly diverse connectivity profiles at the system level –that is, between the S-A axis poles– where sensorimotor systems would be characterized by localized, short-range communication and association systems by long-range projections.

We leveraged connectivity distances to capture changes in functional connectivity profiles by calculating the mean length of the top 10% strongest functional connections for each seed region at the individual level. Crucially, these lengths were computed using geodesic distances on a common cortical template (Schaefer 400 parcellation^61^) –rather than in subject-specific space– in order to decouple developmental shifts in functional connectivity profiles from individual variations in cortical anatomy. By quantifying inter-regional distances on a common template, we ensured that any change in a region’s mean functional connection length reflected shifts in the topological identity of its strongest partners –that is, which regions represent its strongest connections– rather than differences in physical distance stemming from inter-individual anatomical variability or developmental cortical expansion.

Consistent with the other examined features of S-A axis development, we observed significant inter-individual differences in mean functional connection lengths averaged by functional network^65^ (**Fig. 4A**). At the group level, spin-based permutation testing^62^ revealed that these connectivity profiles strongly aligned with the S-A hierarchy (*r* = 0.77, *p*_spin_ < .001), whereby sensorimotor regions were characterized by short-range connections, while association regions favored long-range projections (**Fig. 4B**), consistent with previous findings^67–69^. Across timepoints, we again observed a descriptive trend of systematic polarization of these connectivity profiles by functional network across timepoints, mirroring patterns of S-A axis expansion (**Fig. 4A**). Namely, sensorimotor-leaning networks (visual and somatomotor networks) exhibited global decreases in mean connection length, reflecting an intensification of localized connectivity. Conversely, association-leaning networks (limbic, frontoparietal, and default-mode networks) showed global increases in mean connection length, indicating a shift toward even longer-range functional projections. Transitional networks situated closer to the axis origin, such as the dorsal and ventral attention networks, exhibited less clearly delineated trajectories.

**Figure 4.**
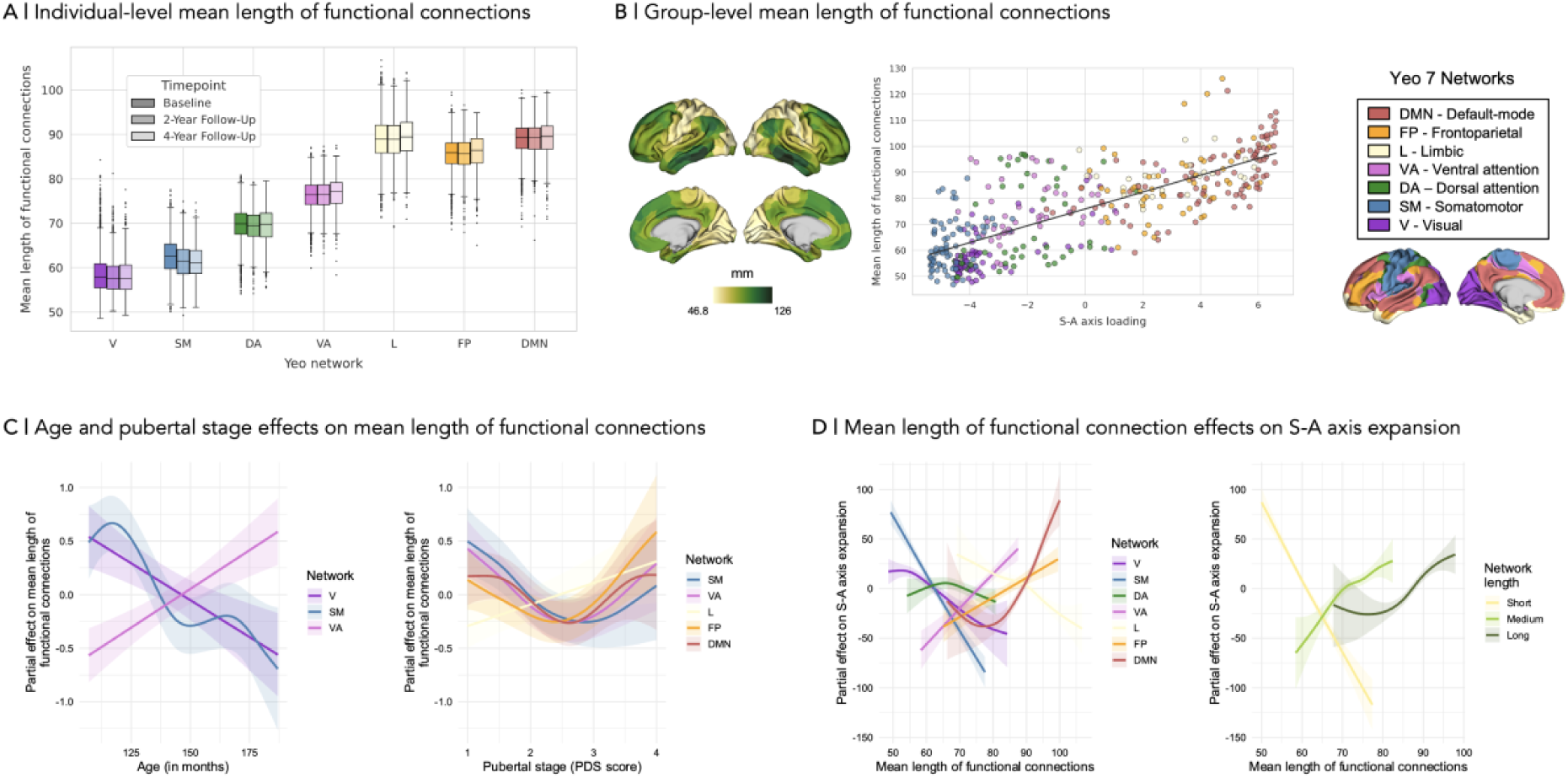
Developmental refinement of functional connectivity profiles characterizing sensorimotor-association (S-A) axis expansion. **A |** Distribution by timepoint and Yeo network of individual-level mean length of functional connections, computed by averaging geodesic distances (on a standard Schaefer 400^61^ template surface) between each seed region and its 10% maximally functionally connected regions, further averaged across regions belonging to each of the seven Yeo networks; **B |** Group-level mean length of functional connections, computed by averaging regional individual-level mean length of top 10% functional connections per seed region across all subjects; scatterplot displays patterns of mean length of functional connections as a function of the seed region’s loading on the adult S-A axis^29^, tested by Spearman correlation and corrected for spatial autocorrelation using spin permutation testing^62^, *r* = 0.77, *p*_spin_ < .001, color-coded by Yeo network; **C |** Estimated partial effects of age and pubertal stage on the mean length of functional connections across Yeo networks (only statistically significant effects are displayed); **D |** Estimated partial effects of mean length of functional connections on S-A axis expansion; Effects are estimated and visualized by individual Yeo network and by aggregating Yeo networks into categories based on their characteristic network length, i.e., short- (V, SM), medium- (DA, VA), and long- (L, FP, DMN) range networks, then averaging connection lengths within each category (all displayed effects are statistically significant). Boxplots are represented by boxes extending from the first to the third quartiles of the data with a line representing the median, whiskers extending from the boxes to the farthest data points within 1.5 times the interquartile range, and fliers representing data points past the ends of the whiskers. Statistical effects were computed with Big Additive Models (BAMs) including fixed effects of chronological age, pubertal stage, sex and total surface area, and random nested effects (subject, family, site). Lines represent the zero-centered component smooths extracted from the BAM models, illustrating the isolated contribution of each predictor to the response while accounting for all other model terms. Shaded ribbons denote 95% Bayesian credible intervals derived from the posterior distribution of the model coefficients, representing the localized uncertainty of the smooth estimate. V, visual; SM, somatomotor; DA, dorsal attention; VA, ventral attention; L, limbic; FP, frontoparietal; DMN, default-mode network.

We again observed distinct effects of chronological age and pubertal stage, here on developmental shifts in functional connectivity profiles (full models explaining 47.0–73.7% of total deviance – detailed test statistics are provided in Supplementary Table 1). Age-dependent changes in connectivity profiles were primarily found in sensorimotor networks, characterized by both linear and non-linear decreases in mean connection length, as well as in the ventral attentional network, characterized by a near-linear increase in mean connection length (**Fig. 4C, left**). In contrast, pubertal stage was related to a more widespread refinement of connectivity profiles additionally involving higher-order association systems, altogether including the somatomotor, ventral attention, limbic, frontoparietal, and default-mode networks (**Fig. 4C, right**). These pubertal trajectories followed a U-shaped profile with an inflection point around mid-puberty. The limbic network was an exception to this trend, instead displaying a near-linear increase in mean connection length across pubertal development.

Finally, we evaluated whether these connectivity shifts were directly related to macroscale S-A axis expansion. Changes in connectivity profiles across all seven networks were indeed significant predictors of S-A axis expansion above and beyond variance already explained by chronological age and pubertal stage (full model accounting for 51.1 % of the total deviance (**Fig. 4D, left**) – detailed test statistics are provided in Supplementary Table 2). To synthesize these network-level effects into more interpretable developmental trends, we aggregated the functional networks into three categories based on their characteristic connection ranges: short-range (visual and somatomotor networks), medium-range (dorsal and ventral attention networks), and long-range (limbic, frontoparietal, and default-mode networks), averaging connection lengths within each category. This analysis revealed a relationship between shifts in connectivity profiles and S-A axis expansion distinctly depending on characteristic connection ranges (**Fig. 4D, right** – detailed test statistics are provided in Supplementary Table 2). Short-range networks exhibited a near-linear negative association with S-A axis expansion, such that the further shortening of their functional connections –representing a relative increase in functional connectivity with regions that are physically nearer– was associated with increasing S-A axis expansion. Conversely, medium- to long-range networks showed positive, non-linear associations with S-A axis expansion, such that the further elongation of their functional connections –representing a relative increase in functional connectivity with regions that are physically more distant– was associated with increasing S-A axis expansion. Sensitivity analyses, in which we computed the categories of connection ranges at the regional rather than network level, confirmed that these patterns were robust to methodological decisions and network definitions, actually showing more linear and distinct trajectories when computed at the regional level (Supplementary Fig. 6).

Collectively, these findings demonstrate that the expansion of the S-A axis, driven by the system-level segregation and differentiation of functional networks, is fundamentally characterized by a polarization of increasingly specialized connectivity profiles, namely the contraction of short-range sensorimotor connections and the elongation of medium-to-long-range projections made by more association-leaning networks. This process represents the maturation of a key feature of the adult functional hierarchy, whereby networks closer to the sensorimotor pole are characterized by their local integration, while networks closer to the association pole are characterized by their distributed communication^67–69^. Importantly, these findings reveal that the specialization of functional connectivity profiles is ongoing across all major functional networks, underscoring that meaningful functional refinement during adolescence is not confined to association systems.

## Discussion

Adolescence is a time of neurodevelopmental reorganization, marked by the global refinement of brain structure and function. Despite puberty’s role as a key driver of biological change during this sensitive period and a substantial source of inter-individual variability in developmental timing, the maturation of the functional cortical hierarchy –represented by the S-A axis, and reflecting a major organizational and developmental principle of functional connectivity– has been solely described as a function of age. Individualized models are also scarce and, where attempted, have relied on alignment to a common reference, introducing systematic biases that distort individual developmental trajectories. Here, we address these gaps by deriving alignment-free continuous and discrete features of S-A axis development at the individual longitudinal level, disentangling the independent contributions of chronological age and pubertal stage on the maturation of the functional hierarchy in a unified explanatory framework. Our findings first reveal substantial heterogeneity across individuals, as well as striking divergence between group- and individual-level characterizations of S-A axis maturation. This highlights that averaging across individuals not only obscures individual differences in developmental trajectories but may misrepresent the direction of developmental trends as well as the normative timing of a developmental milestone for cortical reorganization, namely when the S-A axis becomes the principal gradient explaining the most variance in functional connectivity. We then demonstrate that chronological age and pubertal stage make distinct, additive contributions to the developmental trajectories of different low-dimensional features of hierarchical functional organization. We observe that these developmental effects are related to coordinated shifts in functional network topology that drive the polarization of the functional hierarchy, illustrated by an expansion of the S-A axis representing the global segregation of networks from a systems perspective. At the intrinsic functional level, we show that S-A axis expansion reflects an ongoing specialization of functional connectivity profiles across all major functional networks, highlighting that meaningful functional refinement during adolescence involves sensorimotor and association systems alike. Together, these findings underscore how individual-level approaches are necessary to accurately capture variability in the pace of adolescent cortical maturation, shaped by both chronological age and pubertal development.

Charting neurodevelopmental trajectories across the lifespan is a recent endeavor aiming to describe normative processes of change^34,40,57,64^. While establishing an average healthy population reference is fundamental for our understanding and characterization of pathology as deviations from the norm, this goal tends to shift the focus away from normative individual differences and may underestimate the developmental variance underpinning the population reference by emphasizing average trends in cross-sectional data^70–72^. In addition, the study of individual differences in gradients of brain organization has been further limited by the methodological challenge of relying on orthogonal alignment techniques to allow inter-individual comparability, which comes at the cost of inherently distorting individual variability towards a shared reference^43^. An illustration of this is a recent study charting S-A axis development across the lifespan and using a clustering approach to identify three major phases in the continuous refinement of the S-A axis: initiation (third trimester to perinatal period), establishment (infancy to early childhood), and expansion-stabilization (late childhood to adulthood)^40^. This exemplifies how the use of alignment entirely masked the discrete transition towards adult-like patterns –represented by the S-A axis becoming the principal gradient of functional organization^39,40^– thereby overlooking a key milestone of S-A axis development occurring during adolescence. In the present study, we focused on depicting and preserving individual differences to characterize the developmental trajectories of both continuous and discrete features of functional S-A axis refinement in a longitudinal cohort with up to three timepoints per subject. By computing S-A axis developmental features that did not require any form of alignment –thus preserving idiosyncrasies– our findings revealed substantial heterogeneity in the maturation of the functional hierarchy, with important implications. First, such a high degree of inter-individual variability –in both the continuous refinement of spatial S-A axis patterns and the timing of a key discrete developmental milestone for functional hierarchical development– suggests that maturational trajectories of S-A axis development cannot be meaningfully summarized by group averages. Second, our findings reveal a striking divergence between group- and individual-level characterizations of both our continuous and discrete markers of S-A axis maturation. Whereas the similarity to the adult S-A axis showed a global upward trend at the individual-level across timepoints –confirmed by statistical findings of a positive association with both chronological age and pubertal stage– it paradoxically showed a slightly decreasing trend at the group-level. Group averages also failed to reflect the actual proportion of subjects for whom the discrete transition to adult-like gradient organization had occurred. Specifically, more than half the subjects at baseline (aged 9-11 years) already displayed the S-A axis as the gradient explaining the most variance in functional connectivity, a marker of adult organization. Yet, at the mean group-level, we show that the gradient flip had still not occurred at the 4-year follow-up (including individuals spanning ages 13-16). Previous work only described the gradient flip at the group level, occurring at around ages 11-12^39^ and 14-16^40^, depending on the study – likely reflecting differences in sample composition across studies. Here, with the majority of subjects presenting adult-like gradient organization before it is reflected in the group average, and with opposing trends of similarity to the adult S-A axis at the group and individual levels, our findings point to discrepancies consistent with the ecological fallacy cautioned against in epidemiological research^73^. This pitfall highlights the methodological shortcomings of cross-sectional snapshots in providing representative depictions of individual trajectories. This is in line with evidence of differences between age-related estimates of change produced at the group and longitudinal individual levels^74^, which may be related to individual differences outweighing variance related to age^75^. Our findings therefore nuance previous group-level reports providing a defined average estimate of the age window in which the S-A axis becomes the principal gradient of functional cortical organization^39,40,44^. Instead, we show that measuring individual differences is not only important to characterize inter-individual heterogeneity but also necessary to accurately represent developmental trajectories of cortical reorganization and, in turn, identify the biological and environmental factors that drive them.

Given the collinearity of chronological age and pubertal stage, teasing apart these effects indeed requires leveraging differences in pubertal timing, which are both substantial across individuals and systematic between the sexes – with females experiencing the onset of puberty on average 1-1.5 years earlier than their male counterparts^15^. By largely modeling developmental trajectories as a function of age, the study of macroscale functional organization has been conflating sources of variability, considerably overlooking the role of puberty as a key driver of biological changes during adolescence. Here, we first showed that chronological age and pubertal stage were only moderately correlated, confirming sufficient independent variance beyond their collinearity for models to reliably estimate their unique effects, while highlighting substantial individual differences in pubertal timing. This allowed us to then identify distinct and additive contributions of chronological age and pubertal stage on different markers of S-A axis development. Both factors explained changes in the similarity to the adult S-A axis, while the probability of observing the gradient flip was solely predicted by pubertal stage. Across analyses, the mid-pubertal stage presented as a window of accelerated maturation and reorganization. This may be explained by the non-linear nature of pubertal development itself^76^ and is consistent with previous evidence of greater inter-individual variability in brain development during mid-puberty^71^. Effects of both age and pubertal stage consistently followed shared trajectories across sexes, suggesting that developmental processes pertaining to the refinement of the functional hierarchy may be independent of sex, despite systematic sex differences in the timing and biological consequences of puberty. The only clear sex differences we observed were in females showing consistently greater similarity to the adult S-A axis and greater S-A axis expansion on average. Given previous work reporting that the increasing range of S-A axis loadings during development remains greater in females throughout the lifespan^40,57^, the sex differences we report are more likely to reflect an organizational difference that is conserved throughout the lifespan rather than a fundamental difference in developmental trajectories characterized by anticipated female maturation going beyond earlier pubertal onset. Altogether, our findings align with observations of greater similarities than differences across sexes in the adolescent developing brain^77^. Yet, considering evidence of sex-specific hormonally-modulated neurodevelopment in animal models^78^, as well as increasing anatomical^79,80^ and functional^79^ sex differences during adolescence in humans, further research longitudinally capturing adolescence through early adulthood is needed to examine the progression of sex differences in the functional hierarchy and how they relate to established sex differences in the adult S-A axis^37^.

By examining two key complementary markers of the transition to adult-like gradient organization –the similarity to the adult S-A axis and the gradient flip– our findings more broadly highlight the value of studying different developmental features of functional organization and the importance of separately assessing their respective biological determinants. This consideration is fundamental when comparing our results to findings from previous work investigating different markers of functional development. In fact, while pubertal associations with functional network connectivity have been recently reported^24,25^, others found that heterochronous changes in intrinsic functional activity follow sensorimotor-to-association patterns throughout adolescence as a function of age but not pubertal stage^35^. It is important to note that, in the latter study, the S-A axis was leveraged as a spatiotemporal axis describing patterns in the timing of regional maturation, which is a substantially different endeavor to using the gradient-derived S-A axis as a spatial embedding, as done in the present study. While both approaches pertain to an overarching framework highlighting S-A differentiation as a hierarchical principle of brain organization, one focuses on characterizing patterns of developmental change following a region’s position on the S-A axis, while the other describes the spatial arrangement of regions along a continuum based on their patterns of functional connectivity, characterizing the S-A axis itself. These perspectives are therefore complementary and may –as findings suggest– reveal different developmental processes and biological dependencies, including in their associations to age and puberty.

Our analyses highlight a further distinction in results derived from the use of the S-A axis as a spatiotemporal contextualization as opposed to a spatial embedding. While we replicate trends of age-dependent spatiotemporal changes in functional connectivity strength broadly following the S-A axis^33,34^, we also find that these changes are not associated with S-A axis expansion. Specifically, connectivity strength does not become polarized across sensorimotor and association systems – that is, it does not show opposing developmental trajectories in sensorimotor versus association systems as would be expected if these changes were driving S-A axis expansion. This is consistent with evidence indicating that the S-A axis as a spatial embedding captures changes in organizational features beyond functional connectivity strength^37,57^. In fact, our measure of topographical expansion captures age- and pubertal stage-dependent coordinated shifts in functional networks along the S-A axis towards their respective poles, characterizing the increased differentiation of the S-A axis as a spatial embedding at the system level and extending previous reports of developmental increases in S-A axis range^40,41,57^. We find that S-A axis expansion is instead associated with the system-level functional specialization of connectivity profiles, namely the contraction of short-range connections belonging to sensorimotor networks and the elongation of medium-to-long-range connections of more association-leaning networks. As such, our findings are in line with previous evidence suggesting that the S-A axis as a spatial embedding reflects variation in regional functional connectivity profiles, more specifically the topological identity of a region’s strongest functional partners^37^. Interestingly, we demonstrate that the increasing differentiation of functional connectivity profiles reflects an ongoing functional specialization^56^, involving networks spanning the entire functional hierarchy, thus illustrating that sensorimotor and association systems alike are still maturing at ages 9 to 16. These insights underscore that the established heterochronicity of functional refinement along the spatiotemporal S-A axis –with primary sensorimotor regions maturing before higher-order association regions– should be interpreted in terms of maturational peaks^39,40^ as opposed to the closing of tightly defined critical windows^13^. Adolescence is indisputably a period of heightened development for association systems, marked by a shift from a spatially localized to a globally distributed organization^49^ that is enabled by association regions making long-reaching projections^29,81^. Yet, our findings complement this perspective by showing that, while networks at the association pole indeed extend their long-range communication, those at the sensorimotor pole simultaneously also refine their local integration, *together* driving the expansion of the S-A axis through an increasing polarization of the functional hierarchy. This highlights that the ongoing specialization of sensorimotor regions may be equally relevant to that of association regions for the refinement of the S-A axis specifically.

Beyond the novel insights gained through our study, some limitations should be noted and future directions highlighted. First, to capture the continuous refinement of functional hierarchical development, we computed a measure of similarity to the adult S-A axis by calculating subject-level correlations to a selected adult template^29^ derived from the Human Connectome Project Young Adults dataset^82^. We cannot exclude that differences reflected in our measure of adult-like similarity may partly stem from differences between the youth and adult samples and their measurements having been obtained from different neuroimaging collection and analysis pipelines. While, this discrepancy is inevitable regardless of the chosen adult reference sample, the high correlations reached by our measure of adult-like similarity (ranging *r* = 0.37-0.95) confirm the overall comparability of the youth and adult samples, while the metric’s large range of variability further demonstrates its sensitivity to individual differences. We intentionally selected the original adult S-A axis template^29^ as it is openly available and commonly used as a reference, thus facilitating reproducibility of findings. Importantly, the sample from which it was derived is also based in the United States and includes young adults, which is essential to avoid confounding effects of ageing characterized by patterns of S-A axis contraction^57^ and decreased global network segregation^34,56,83^ in older age, as they are opposite to developmental trends observed during adolescence. Secondly, in our comparison of chronological age and pubertal stage effects, the estimation of pubertal development has a measurement error, whereas age does not. We selected parental reports of pubertal development given their greater reliability relative to child reports in this age range^84–86^, but future data collection efforts should consider Tanner staging by a trained healthcare professional, as it is the gold medical standard^87^. Furthermore, while it is the endocrine changes associated with puberty that are expected to drive neurodevelopmental mechanisms, our consideration of pubertal stage solely based on physical characteristics only acts as a proxy for hormonal effects. However, evidence from the ABCD sample demonstrates significant correlations between steroid hormone levels and pubertal stage, suggesting that the latter may also serve as an estimate of pubertal neuroendocrine changes^88^. Moreover, our primary aim was to disentangle chronological age and pubertal stage contributions to functional brain development. Given that physical pubertal development reflects cumulative hormonal changes over time, whereas steroid hormone levels fluctuate within subject, show substantial inter-individual variability^89^, and are prone to measurement error (including ABCD study’s lack of standardization in the time of hormonal data collection^90^), physical pubertal changes represent a more suitable and reliable marker of neuroendocrine influences than single hormone measurements for the scope of this study. Future research using animal models for tighter experimental control and an assessment of neuroendocrine effects via more invasive measurements would allow to further disentangle age from pubertal effects at a more mechanistic hormonal level. Thirdly, our sample ranges ages 9 to 16, thereby not providing complete coverage of the adolescent developmental period. Although the three timepoints included in this study comprised subjects together capturing the full span from early to late puberty, future timepoints of the ongoing ABCD study data collection will provide greater statistical power at later pubertal stages, especially in males. This will also allow to investigate the effects of pubertal tempo, describing the pace of pubertal development, which has also been shown to influence brain development^91^. Future work further dissecting pubertal effects on neurodevelopment should additionally consider biological and environmental determinants of pubertal timing and tempo –such as (epi)genetic mechanisms^92^– given their consequential effects on mental health^93,94^. It is only by characterizing inter-individual differences in adolescent brain development, and by understanding their normative sources of variability, that we will be in a position to identify –and address– risk factors for psychopathology.

## Methods

### Participants

Our sample comprised of data from the Adolescent Brain Cognitive Development (ABCD) study^58^, the largest longitudinal consortium study of brain development in the United States aiming to capture wide-ranging factors affecting adolescent neurodevelopment and health. Participants were recruited across 21 research sites with the aims of producing a cohort that is representative of the sociodemographic heterogeneity and overall diversity of the United States population. Detailed recruitment considerations and sampling procedures are described elsewhere^95^.

The present study included demographic, pubertal development, and neuroimaging data of *N*_observations_ = 6323 (including 4919 unique subjects and 4409 unique families) in a longitudinal design tracking functional cortical maturation across three waves of data collection: baseline (107-132 months; *n* = 4064), 2-year (129-165 months; *n* = 1296), and 4-year (151-187 months; *n* = 963) follow-ups (total age range: 8.9-15.6 years, with a mean age of 11.0 ± 1.7 years (132.1 ± 19.9 months), and 3220 females against 3103 males across observations). Parent-reported race distribution amongst unique subjects was 70.8 % White, 11.7 % mixed race, 10.3 % Black, 1.9 % Asian, 0.6 % Native American, 0.1 % Pacific Islander, 3.5 % other race, and 1.1 % not reported. All subjects gave their verbal assent –and their parents provided written consent–prior to study participation. Data was obtained from the National Institute of Mental Health (NIMH) Data Archive (Release 5.1), and inclusion of subjects was based on local availability of data and quality control criteria (detailed information in the Supplementary Methods, Supplementary Fig. 7).

### Pubertal development

Given our focus on the biological process of puberty, we used parent-reported sex assigned at birth in our analyses to account for biological sex.

The Pubertal Development Scale (PDS), a standardized questionnaire assessing physical pubertal development, was used to characterize pubertal stage^63^. It includes five items in total, assessing perceived changes in body hair, skin and height (across sexes), as well as voice deepening and facial hair growth (in males), and breast development and menarche (in females). All items were scored on four-point Likert scales, with higher scores indicating more advanced pubertal development. For the menarche item, a score of one indicates “menstruation has not yet begun” and four indicates “menstruation has begun”. PDS scores were computed by averaging item scores, yielding a final score ranging from one (no physical signs of pubertal development) to four (completed pubertal development). We considered subjects with available data for at least three of five items. The ABCD administers the PDS questionnaire to both youth and parents, but we only considered parental reports given their greater reported reliability, particularly in late childhood and early adolescence^84–86^.

### MRI data acquisition

The ABCD’s MRI data was acquired by following harmonized imaging protocols across three 3T scanner platforms (Philips, Siemens Prisma, and General Electric 750) with standard adult-size multi-channel coils suitable for multiband echo planar imaging (EPI) acquisitions. MRI data was collected in the axial plane. Structural images were acquired using a T1-weighted magnetization-prepared rapid acquisition gradient echo sequence and a T2-weighted fast spin echo with a variable flip angle sequence, both at 1 mm isotropic voxel resolution. Functional images were acquired in four separate resting-state runs of 5 minutes each, using simultaneous multi-slice/multiband EPI with fast integrated distortion correction and the following parameters: 2.4 mm isotropic voxel resolution, matrix = 90×90, 60 slices, FOV = 216×216, TR = 800 ms, TE = 30 ms, flip angle = 52°). More detail and information on image acquisition, sequence parameters and decisions is reported elsewhere^96^.

### MRI data processing

MRI data processing was restricted to participants meeting the ABCD study’s curated quality control criteria for T1-weighted and resting-state fMRI acquisitions^97^. The preprocessing scripts described below are available in our GitHub repository.

### Structural MRI

T1w and T2w images were preprocessed with FreeSurfer (v7.2.0; https://surfer.nmr.mgh.harvard.edu/), using the recon-all command which included image registration to standard space, tissue segmentation, and initial cortical surface reconstruction^98^. Notably, the T2-weighted acquisition was utilized to refine the pial surface boundary^99^.

In previous work, we show that total surface area is the measure of brain size most strongly associated with the S-A axis (relative to intracranial volume and total brain volume)^37^. Therefore, we computed total surface area by extracting surface data from native-space FreeSurfer surfaces by hemisphere and summing the extracted surface data within and between hemispheres. We then included this measure of brain size as a covariate in all of our statistical models, in line with standard practice in developmental neuroimaging and when examining sex differences.

### Functional MRI

Resting-state functional MRI images were preprocessed with fMRIPrep (v22.1.1; https://fmriprep.org/en/stable/)^100^. The pipeline used the precomputed cortical reconstructions and brain segmentations generated by FreeSurfer. Functional images were co-registered to the T1-weighted anatomical template using boundary-based registration^101^. Preprocessed data were resampled into several output spaces, including native functional space and the fsaverage surface template (10k density). Additionally, grayordinate files were generated at 91k resolution (containing 32k vertices per hemisphere) to project BOLD timeseries onto the fsLR cortical surface^102^.

Finally, we used the XCP-D pipeline (v0.10.1, https://xcp-d.readthedocs.io/en/latest/) for the postprocessing of the functional resting-state fMRIPrep output, aimed at mitigating motion-related noise and artifacts^103^. This pipeline treated each run separately and included the removal of dummy volumes, despiking, mean-centering, linear detrending, bandpass filtering (0.01–0.08 Hz) and spatial smoothing (6 mm FWHM) of the BOLD signal. Despiking consists in the temporal censoring of high-motion outlier volumes^104^, which were detected using framewise displacement with a threshold of 0.3mm and a head radius of 50mm^105^. Detrending was performed with linear regression by regressing 20 nuisance regressors from the BOLD data. The custom nuisance regressors were estimated based on the fMRIPrep preprocessed timeseries output and included six motion parameters (xyz translations, xyz rotations), their temporal derivatives and squared terms, as well as mean signal from cerebrospinal fluid and white matter. Runs with less than 240 seconds of usable data after removal of dummy and high motion volumes were not processed further. For each run, XCP-D extracted processed functional timeseries using Connectome Workbench^102^ for the ‘4S456Parcels’ atlas (corresponding to 400 Schaefer cortical parcels^61^ and 56 subcortical parcels^102,106–108^), concatenated timeseries, and directly outputted functional connectivity matrices reflecting pairwise Pearson correlations. We then manually excluded subcortical parcels, averaged across the outputted matrices for subjects who had a minimum of 2 runs, and conducted a Fisher *r*-to-*z* transformation of the matrices, yielding a final 400×400 normalized connectivity matrix per subject consisting of a minimum of 8 minutes of total scan time across runs.

### Computation of cortical gradients of brain organization and main features of S-A axis development

This study leveraged cortical gradients of brain organization to study the refinement of the macroscale functional hierarchy across adolescence. To this end, we computed the functional S-A axis and derived a range of features capturing S-A axis development.

The S-A axis can be obtained in a data driven manner by reducing the dimensionality of functional connectivity matrices to yield low dimensional gradients of functional organization^29^. These representations are axes describing the global structure of the functional connectivity matrices, along which cortical regions that show similar connectivity patterns are represented closer together, whereas regions with low functional timeseries covariance are represented farther apart, as indexed by the loadings of the cortical regions on these gradients. Typically, ten gradients of brain organization are computed (G1-G10), ordered by their explained variance. Previous findings consistently report that, in adults, the S-A axis is represented by the primary gradient (G1), explaining the most variance^29,39,44^, whereas in children, it is represented by the secondary gradient (G2), explaining the second most variance^39,44^.

We used the BrainSpace Python toolbox^109^ to compute ten macroscale functional connectivity gradients and yield the S-A axis at both group and individual levels. Group-level gradients were computed per data collection timepoint (i.e., baseline, 2-year and 4-year follow-ups) from mean functional connectivity matrices averaged across study participants within each timepoint. Individual-level gradients were computed from each subject’s mean functional connectivity matrix averaged across scan runs. For consistency with previous studies and to allow for the comparison with the adult S-A axis template^29^, we used the following steps and parameters to compute gradients. Briefly, functional connectivity matrices were transformed into affinity matrices using cosine distance, yielding positive weights between zero and one. Then, diffusion map embedding, a nonlinear manifold learning algorithm, was used to reduce the complex, high-dimensional affinity matrices to low-dimensional representations combining geometry with the probability distribution of data points^110^. Only the top 10% row-wise matrix values were considered for the data reduction (i.e., 90% threshold), representing each seed region’s top 10% maximally functionally connected regions. The α parameter, controlling the influence of the sampling points density on the manifold, was set to 0.5 (where α = 0 (maximal influence) and α = 1 (no influence)), and the t parameter, controlling the scale of eigenvalues, was set to 0. This yielded ten macroscale functional connectivity gradients, from which the S-A axis could be identified. At the mean group level, S-A axis patterns were identified in G2 at all three timepoints. At the individual level, we observed substantial heterogeneity in which gradient represented the S-A axis.

We then derived two novel primary features of S-A axis development at the individual level. First, a continuous measure of similarity to the adult S-A axis, quantified by the absolute Spearman correlation coefficient of the specific gradient that most strongly correlated with the adult group-level mean S-A axis template^29^. Second, the occurrence of the gradient flip, a binary measure quantifying whether S-A patterns were reflected in G1, representative of the transition toward adult-like gradient organization. We also computed a distance to gradient flip metric for each participant as the difference between the scaled variance explained (%) by the S-A axis and the highest-variance non-S-A gradient, with negative values indicating that the gradient flip has not yet occurred and positive values indicating that the S-A axis is the gradient explaining the most variance. In the process of computing these features of S-A axis development, we did not align individual-level gradients to any reference group-level average (e.g., via Procrustes alignment) contrary to common practice, in order to preserve inter-individual differences across subjects. This was possible given that our downstream statistical analyses were conducted on derived summary metrics, for which cross-subject comparability at the level of raw gradient loadings –typically achieved via alignment– is not required. In fact, avoiding alignment ensured that S-A axis loadings were not artificially distorted toward the reference template, thus preserving inter-individual variability in the continuous adult-like similarity. Crucially, it also allowed to capture the discrete gradient flip occurrence at the individual level, which is dependent on the native, unaligned rank of the principal gradients. Because Procrustes alignment typically reorders and rotates individual gradients to maximize their correspondence to the reference template, applying it would force the individual-level gradient ranks to mirror those of the reference template, masking inter-individual differences in the categorical shift of S-A patterns from G2 to G1, thereby obscuring a fundamental feature of cortical reorganization during the adolescent transition. Therefore, by maintaining gradients in their native unaligned space, we ensured that inter-individual differences in both the continuous spatial refinement (similarity to the adult S-A axis) and the discrete hierarchical reordering (gradient flip) were captured.

### Computation of S-A axis expansion

Building on recent centroid-based frameworks for quantifying network dispersion in gradient space^37,38,64^, we developed a novel, individual-level measure of S-A axis expansion to capture topological system-level shifts in functional networks, specifically their divergence from the hierarchical center toward the sensorimotor and association poles.

We first identified network centroids for each of the seven Yeo networks^65^ by computing the median loading across constituent regions, which represents the anchoring position of each network on the S-A axis. S-A axis expansion was then calculated as the sum of squared values of these seven network centroids, which effectively represents the aggregate hierarchical distance of these networks from the S-A axis origin (zero), reflecting the global segregation of networks at the system level. We identified S-A axis expansion outliers as values 1.5 times greater than the third quartile and below the first quartile, and excluded them from further analyses (*N*_subsample_ = 6109).

To confirm the robustness of the S-A axis expansion metric and the validity of our results beyond methodological decisions and network definitions, we conducted sensitivity analyses in which we computed the S-A axis expansion metric at the regional level, namely by calculating the sum of squared values of regional S-A axis loadings instead of network centroids (Supplementary Fig. 3A-C). Additional sensitivity analyses were carried out to confirm that the validity of our results was independent of outlier exclusion (Supplementary Fig. 3D-F).

### Computation of functional connectivity profiles

To quantify developmental shifts in functional connectivity profiles, we estimated the mean length of functional connections per region –namely the mean distance between each seed region and its top 10% strongest functional partners– at the individual level using geodesic distance, representing the shortest path between two regions along the curvature of the cortical mantle.

We first derived a common template geodesic distance matrix following the Micapipe protocol^111,112^. Using the Connectome Workbench^102^, we generated a midthickness cortical surface in fsaverage space by averaging the white matter and pial surfaces. For each of the 400 regions in the Schaefer parcellation^61^, we identified a representative centroid vertex, defined as the vertex with the shortest summed Euclidean distance to all other vertices within that parcel. Then, we applied Dijkstra’s algorithm^113^ to compute the pairwise geodesic distances between all 400 centroid vertices on the midthickness mesh. This resulted in a standardized distance matrix representing physical distances in the cortical Schaefer 400 template space.

For each subject, we identified the top 10% strongest functional connections of every seed region, which also represent the functional connections that underwent dimensionality reduction in the computation of the S-A axis and its developmental features. We then extracted the corresponding geodesic distances for these specific connections from our common template matrix and averaged them to yield regional (seed-wise) measures of mean functional connection lengths. By calculating these inter-regional distances on a common template rather than in subject-specific space, we effectively decoupled topological shifts in functional organization from individual variations in cortical anatomy. Specifically, we ensured that any change in a region’s mean connection length reflected shifts in the topological identity of its strongest partners –that is, which regions represented its strongest connections– rather than differences in physical distances stemming from inter-individual variability or developmental cortical expansion.

We further summarized these regional connectivity profiles at the network-level by averaging across the mean lengths of functional connections of all regions belonging to the same Yeo network^65^. Then, to synthesize these network-level effects into more interpretable developmental trends, we also aggregated the functional networks into three categories based on the characteristic length of their connections: short-range (visual and somatomotor networks), medium-range (dorsal and ventral attention networks), and long-range (limbic, frontoparietal, and default-mode networks), and averaged connection lengths within each category. To confirm the robustness of these categories and the validity of our results beyond methodological decisions and network definitions, we conducted sensitivity analyses in which we defined these categories of characteristic connection ranges at the regional level by assigning each region (rather than network) to one of the three categories (short, medium, long) based on the mean length of its top functional connections and then averaging across distances within each category (Supplementary Fig. 6).

### Statistical analyses

Statistical analyses were conducted in R (v4.5.0). To capture the non-linear developmental trajectories of the S-A axis, we utilized Big Additive Models (BAMs) –a scalable extension of Generalized Additive Models (GAMs) optimized for large-scale datasets– using the mgcv package^114^. These models systematically accounted for the effects of major factors associated with brain development to isolate their relative contributions, namely age, pubertal stage, sex, and total surface area as the measure of brain size most strongly associated with variability in the S-A axis^37^. Furthermore, to account for the nested structure of the ABCD dataset –where subject observations at a given timepoint are nested within families and research sites– we incorporated a multi-level random-effects structure, modelling site, family-by-site, and subject-by-family-by-site as random-effect basis terms (bs=”re”). This approach accounted for the non-independence of errors across timepoint, family and site clusters.

In our models, maturational smoothers were estimated using penalized regression splines. We used fast Restricted Maximum Likelihood (fREML) for smoothness selection, which provides a robust and computationally efficient estimation of the smoothing parameters. To prevent overfitting, we set the basis dimension to k = 10 (default) and employed penalized thin-plate regression splines (fx = FALSE). We also used the select = TRUE argument to allow for null-space penalization, effectively enabling the model to penalize uninformative smooth terms to zero. To ensure that we were not underfitting, namely that the basis dimension was sufficient to represent the degree of non-linearity in the biological signal, we performed post-hoc basis dimension adequacy checks using mgcv’s k.check function for each model. This allowed us to confirm that the Effective Degrees of Freedom (*EDF*; representing the degree of non-linearity where *EDF* = 1 indicates a linear effect and *EDF* > 1 indicates increasing complexity) remained well below k′ (the maximum possible flexibility), and that the k-index and associated *p* values did not indicate underfitting.

To test for sex differences in chronological age and pubertal stage effects, we used a reference-level parameterization with ordered factors. By converting sex into an ordered factor, we decomposed the smooth terms into a reference smooth (representing the main effect of the variable) and a difference smooth (representing sex-specific deviation from the reference). This allowed us to explicitly test the statistical significance of the sex-by-chronological age and sex-by-pubertal stage interactions. To report and visualize marginal effects, we generated posterior predictions from the BAMs by sex. For this, continuous covariates were held at their mean values, and random nested effects (subject, family and site) were excluded in order to produce population-level trajectories, effectively marginalizing over the nested structure of the sample. In the case of effects involving the mean lengths of functional connections by network and by characteristic connection lengths, we plotted zero-centered component smooths extracted from the BAM models, illustrating the isolated contribution of each predictor to the response while accounting for all other model terms. This decision was made for visualization purposes, in order to include all networks and categories of functional connection lengths in the same figure.

### Supplementary analyses

We performed a range of supplementary analyses to ascertain the specificity and robustness of our findings. These included assessing the relationship between our measures of S-A axis maturation (Supplementary Fig. 1) and exploring effects related to changes in functional connectivity strength (Supplementary Fig. 2), as well as sensitivity analyses to confirm that the reported developmental effects were not driven by methodological decisions nor network definitions in the computation of S-A axis expansion (Supplementary Fig. 3), potential confounders such as socioeconomic status (SES; specifically, parental education; Supplementary Fig. 4) and body mass index (BMI; Supplementary Fig. 5), nor methodological decisions in the definition of categories of characteristic connection ranges (Supplementary Fig. 6). These analyses and their results are all described and reported in the Supplementary Materials.

## Data availability

All data needed to evaluate the conclusions of the paper are present in the paper and in the Supplementary Materials, and are further available upon request

Data used in the preparation of this article were obtained from the [Adolescent Brain Cognitive Development^SM^ (ABCD) Study](https://abcdstudy.org/), held in the NIH Brain Development Cohorts Data Sharing Platform. This is a multisite, longitudinal study designed to recruit more than 10,000 children age 9-10 and follow them over 10 years into early adulthood.

The ABCD Study® is supported by the National Institutes of Health and additional federal partners under award numbers U01DA041048, U01DA050989, U01DA051016, U01DA041022, U01DA051018, U01DA051037, U01DA050987, U01DA041174, U01DA041106, U01DA041117, U01DA041028, U01DA041134, U01DA050988, U01DA051039, U01DA041156, U01DA041025, U01DA041120, U01DA051038, U01DA041148, U01DA041093, U01DA041089, U24DA041123, U24DA041147. A full list of supporters is available at [Federal Partners - ABCD Study](https://abcdstudy.org/about/federal-partners/).

The ABCD dataset grows and changes over time. The ABCD data used in this report came from NIMH Data Archive (NDA) Release 5.1 (DOI: 10.15154/z563-zd24). DOIs can be found at https://nda.nih.gov/abcd/abcd-annual-releases for NDA releases and at https://doi.org/10.82525/jy7n-g441 for newer NIH Brain Development Cohorts (NBDC) releases.

## Code availability

Analyses were conducted in Python and R: The code used in this manuscript is available at https://github.com/biancaserio/dev_age_puberty_gradients. The code and tutorials for functional gradient decomposition can further be found at https://brainspace.readthedocs.io/en/latest/index.html.

## Author contributions

Conceptualization: BS, SLV. Data processing: BS, LW, FH. Analysis and visualization: BS. Input on analysis: LD, MM, and SLV. Writing–original draft: BS. Writing–review and editing: BS, LD, MM, LW, FH, DSM, SBE, SLV. Supervision: SLV.

