## Supplementary Materials for "Individual differences reveal distinct age and pubertal contributions to the refinement of the functional cortical hierarchy during adolescence"

**This document includes:**

Supplementary Results
- Main Analyses – Test Statistics (Supplementary Tables 1-2)
- Supplementary Analyses (Supplementary Figures 1-2)
- Sensitivity analyses (Supplementary Figures 3-6)

Supplementary Methods
- Participant inclusion (Figure 7)

### Supplementary Results

#### Main Analyses – Test Statistics

| | *F* | *p* | *EDF* | | *t* | *p* | Deviance explained |
|---|---|---|---|---|---|---|---|
| **Visual** | | | | | | | |
| *Smooth terms* | | | | *Linear terms* | | | 56.4 % |
| Age | 6.42 | < .001* | 0.93 | Sex (M) | 4.48 | < .001* | |
| Pubertal stage | 1.06 | 0.091 | 1.29 | Total SA | 1.25 | 0.213 | |
| **Somatomotor** | | | | | | | |
| *Smooth terms* | | | | *Linear terms* | | | 60.4 % |
| Age | 13.70 | < .001* | 3.94 | Sex (M) | -3.21 | 0.001 | |
| Pubertal stage | 4.37 | 0.002 | 2.35 | Total SA | 4.67 | < .001* | |
| **Dorsal attention** | | | | | | | |
| *Smooth terms* | | | | *Linear terms* | | | 73.7 % |
| Age | 1.17 | 0.575 | 4.20 | Sex (M) | -3.71 | < .001* | |
| Pubertal stage | 3.56 | 0.066 | 2.52 | Total SA | 5.04 | < .001* | |
| **Ventral attention** | | | | | | | |
| *Smooth terms* | | | | *Linear terms* | | | 64.1 % |
| Age | 11.27 | < .001* | 0.96 | Sex (M) | -5.43 | < .001* | |
| Pubertal stage | 6.36 | < .001* | 2.51 | Total SA | 4.93 | < .001* | |
| **Limbic** | | | | | | | |
| *Smooth terms* | | | | *Linear terms* | | | 47.0 % |
| Age | 0.00 | 0.661 | 0.00 | Sex (M) | -2.00 | 0.046* | |
| Pubertal stage | 1.42 | 0.018 | 0.86 | Total SA | -4.12 | < .001* | |
| **Frontoparietal** | | | | | | | |
| *Smooth terms* | | | | *Linear terms* | | | 68.4 % |
| Age | 0.99 | 0.237 | 1.26 | Sex (M) | -3.02 | 0.003* | |
| Pubertal stage | 4.79 | 0.006* | 2.59 | Total SA | 4.40 | < .001* | |
| **Default-mode** | | | | | | | |
| *Smooth terms* | | | | *Linear terms* | | | 64.4 % |
| Age | 1.04 | 0.225 | 1.19 | Sex (M) | -0.20 | 0.840 | |
| Pubertal stage | 3.18 | 0.046* | 3.01 | Total SA | 0.14 | 0.885 | |

**Supplementary Table 1. Independent models testing for age and pubertal stage effects on the mean length of functional connections by Yeo network (results displayed in Fig. 4C).** Statistical effects were computed with Big Additive Models (BAMs) including fixed effects of chronological age, pubertal stage, sex and total surface area (SA), and random nested effects (subject, family, site). * denotes statistical significance at $p < 0.05$.

| | $F$ | $p$ | $EDF$ | | $t$ | $p$ | Deviance explained |
|---|---|---|---|---|---|---|---|
| **Yeo networks** | | | | | | | |
| *Smooth terms* | | | | *Linear terms* | | | 51.1 % |
| Visual | 17.28 | < .001* | 2.81 | Sex (M) | -7.42 | < .001* | |
| Somatomotor | 54.33 | < .001* | 1.44 | Total SA | 0.92 | 0.359 | |
| Dorsal attention | 1.99 | 0.003* | 2.18 | | | | |
| Ventral attention | 13.71 | < .001* | 1.23 | | | | |
| Limbic | 13.10 | < .001* | 3.25 | | | | |
| Frontoparietal | 4.75 | < .001* | 0.96 | | | | |
| Default-mode | 47.89 | < .001* | 3.83 | | | | |
| Age | 1.06 | 0.006* | 0.87 | | | | |
| Pubertal stage | 1.45 | < .001* | 0.92 | | | | |
| **Connection length categories (network-based)** | | | | | | | |
| *Smooth terms* | | | | *Linear terms* | | | 49.3 % |
| Short | 76.43 | < .001* | 1.59 | Sex (M) | -6.25 | < .001* | |
| Medium | 7.12 | < .001* | 3.25 | Total SA | 1.15 | 0.250 | |
| Long | 17.58 | < .001* | 2.97 | | | | |
| Age | 2.24 | < .001* | 0.92 | | | | |
| Pubertal stage | 1.06 | 0.004* | 0.89 | | | | |
| **Connection length categories (region-based)** | | | | | | | |
| *Smooth terms* | | | | *Linear terms* | | | 50.3% |
| Short | 34.32 | < .001* | 2.61 | Sex (M) | -7.27 | < .001* | |
| Medium | 4.63 | 0.003 | 2.62 | Total SA | 2.17 | 0.030 | |
| Long | 11.88 | < .001* | 2.26 | | | | |
| Age | 4.54 | < .001* | 0.96 | | | | |
| Pubertal stage | 1.37 | 0.003 | 0.91 | | | | |

**Supplementary Table 2. Independent models testing for mean length of functional connection effects on sensorimotor-association (S-A) axis expansion, by Yeo network, network-based connection length categories (results displayed in Fig. 4D), and region-based connection length categories (results displayed in Supplementary Fig. 6).** Statistical effects were computed with Big Additive Models (BAMs) including fixed effects of chronological age, pubertal stage, sex and total surface area (SA), and random nested effects (subject, family, site). * denotes statistical significance at $p < 0.05$.

#### Supplementary Analyses

In order to better understand the relationship between the two novel features of S-A axis development under study, i.e., the similarity to the adult S-A axis and the occurrence of the gradient flip, we fitted two models testing for the effects of individual similarity to the adult template on the probability of observing the gradient flip, both when including and not including age and pubertal stage as covariates in our model (Supplementary Fig. 1A-B). In both cases, adult-like similarity was significantly associated with the gradient flip probability, suggesting that this relationship holds above and beyond the variance explained by chronological age and pubertal stage.

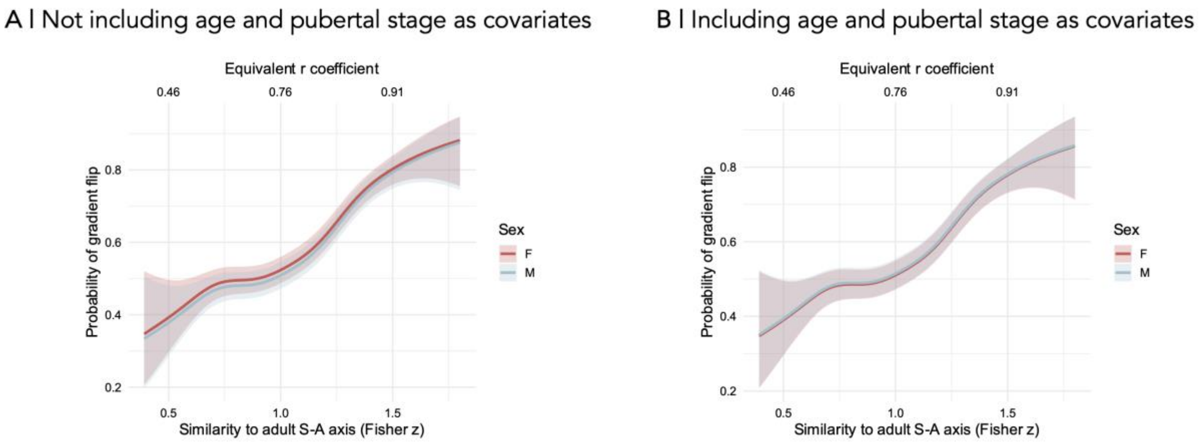


**Supplementary Figure 1. Relationship between similarity to the adult S-A axis and the occurrence of the gradient flip. A |** Predicted marginal trajectories for similarity to adult S-A axis effect on gradient flip by sex, not including chronological age and pubertal stage as covariates in the model; **B |** Predicted marginal trajectories for similarity to adult S-A axis effect on gradient flip by sex, including chronological age and pubertal stage as covariates in the model. Statistical effects were computed with Big Additive Models (BAMs) including fixed effects of sex and total surface area (as well as age and pubertal stage in B), and random nested effects (subject, family, site). Predictions were generated by holding all continuous covariates at their mean values and by excluding random nested effects to produce population-level trajectories. Shaded ribbons represent 95% Bayesian credible intervals derived from the posterior distribution of the model coefficients, representing the uncertainty in the population-level response by excluding the variance attributed to the random nested effects of subject, family, and site.

Previous work shows that the age-dependent development of functional connectivity strength is systematically aligned with the S-A axis, whereby functional connectivity strength increases in primary sensorimotor regions and decreases in higher order association regions from childhood to early adulthood (Luo et al. 2024; Sun et al. 2025). Although evidence also suggests that changing patterns of functional connectivity strength are not reflected in shifts in S-A axis loadings (i.e., when the functional S-A axis is characterized as a spatial embedding) (Serio et al. 2024; Taylor et al. 2026), we still explored whether changes in the strength of functional connections might be related to S-A axis expansion during development.

To allow comparisons with previous findings (Luo et al. 2024; Sun et al. 2025), we computed functional connectivity strength at both the network and regional levels. At the network level, we averaged functional connectivity strength between each seed region and its 10% maximally functionally connected regions at the individual level, further averaged across regions belonging to each of the seven Yeo networks. At the regional level, regions were equally divided into 10 bins based on their position on the adult S-A axis template (Margulies et al. 2016) and functional connectivity strength was further averaged within each bin at the individual level.

Consistent with the other features of S-A axis development under study, we observed significant inter-individual differences in mean functional connection lengths averaged by functional network (Yeo et al. 2011) (Supplementary Fig. 2A). At the group level, spin-based permutation testing (Alexander-Bloch et al. 2018) revealed that these connectivity profiles were negatively associated with the S-A axis hierarchy ($r$ = -0.30, $p_{\text{spin}}$ = 0.010), with sensorimotor networks characterized by stronger mean functional connectivity strength than association networks (Supplementary Fig. 2B). Across timepoints, we again observed a descriptive trend of mean connectivity strength polarization by functional network, hinting at patterns of S-A axis expansion (Supplementary Fig. 2A). Namely, sensorimotor-leaning networks (visual and somatomotor networks) exhibited global increases in mean connection strength with time, while association-leaning networks (limbic, frontoparietal, and default-mode networks) showed global decreases in mean connection strength, in line with previous findings (Luo et al. 2024; Sun et al. 2025). Transitional networks situated near the axis origin, such as the dorsal and ventral attention networks, exhibited less distinct trajectories.

Contrary to findings on the developmental refinement of functional connectivity profiles, statistical modeling of the developmental shifts in functional connectivity strength did not reveal robust effects of chronological age and pubertal stage (full models explaining 54.8–65.0% (network-level) and 54.1-63.9% (region-level) of total deviance). In fact, age-dependent changes in functional connectivity strength were only statistically significant in the limbic network (Supplementary Fig. 2C, bottom left) and in bins 1, 2 and 7 (Supplementary Fig. 2C, bottom right). Nevertheless, at the regional level, general trajectories of effects –albeit largely lacking statistical significance– were in line with previous reports of increasing functional connectivity strength in sensorimotor regions and decreasing functional connectivity strength in associations regions as a function of age (Supplementary Fig. 2C, top right) (Luo et al. 2024; Sun et al. 2025). Pubertal stage-dependent changes in functional connectivity strength were also only statistically significant in a few instances, namely in the frontoparietal network (Supplementary Fig. 2C, bottom left) and in bins 1 and 5 (Supplementary Fig. 2C, bottom right).

Finally, we evaluated whether these connectivity shifts were directly related to macroscale S-A axis expansion. At the network level, an integrated model demonstrated that changes in connectivity strength across all but the visual network were significant predictors of S-A axis expansion above and beyond variance already explained by age and puberty (full model accounting for 53.5 % of the total deviance, Supplementary Fig. 2D, left). However, unlike effects of functional connectivity profiles on S-A axis expansion, functional connectivity strength effects did not show systematic patterns across the cortical hierarchy. For instance, while S-A axis expansion displayed a positive relationship with functional connectivity strength in the default mode network, this relationship was negative with functional connectivity strength in the frontoparietal network – despite both networks being characterized as association systems. Furthermore, a rather flat, slightly negative relationship was observed in the functional connectivity strength of sensorimotor systems, such as the visual and somatomotor networks. At the regional level, findings were similar. An integrated model demonstrated that changes in connectivity strength across all bins were significant predictors of S-A axis expansion above and beyond variance already explained by age and puberty, although the age effect was no longer statistically significant in this model (full model accounting for 60.7 % of the total deviance, Supplementary Fig. 2D, right). Yet, functional connectivity strength effects again did not show systematic patterns across the cortical hierarchy, with similar contrasting directions of effects across systems.

Collectively, these findings replicate effects of functional connectivity strength development aligning with the S-A axis as a function of age (Luo et al. 2024; Sun et al. 2025), supporting the characterization of the S-A axis as a spatiotemporal axis of maturation. Nevertheless, they demonstrate that developmental changes in functional connectivity strength do *not* show systematic and characteristic effects on S-A axis expansion that are reminiscent of a polarization between sensorimotor and association systems – as is the case with effects of maturing functional connectivity profiles. Therefore, in line with previous evidence of a lack of correspondence between the loadings of the S-A axis as a spatial embedding and functional connectivity strength (Serio et al. 2024; Taylor et al. 2026), S-A axis expansion does not appear to reflect a polarization of functional connectivity strength patterns.

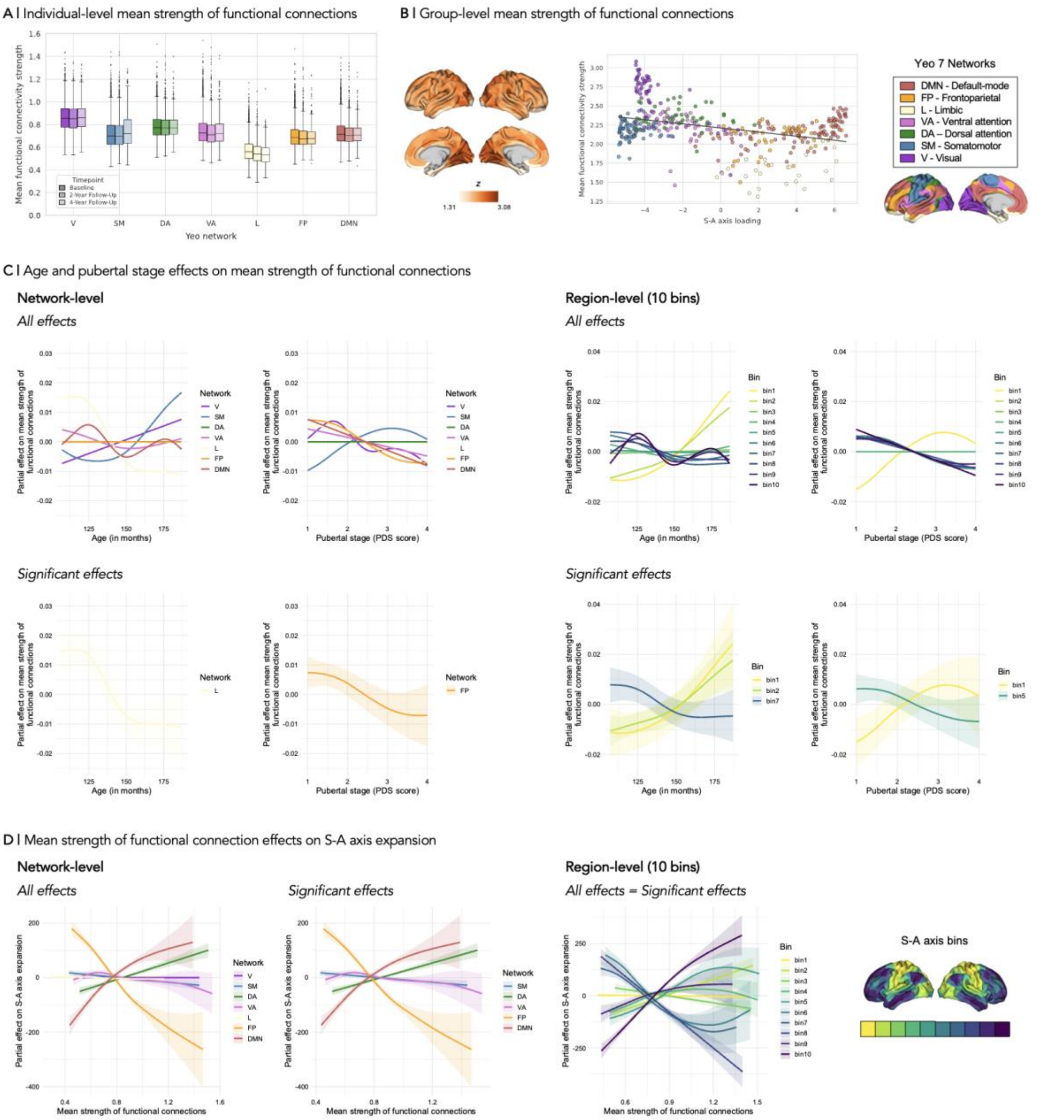


**Supplementary Figure 2. Associations between the developmental refinement of functional connectivity strength and sensorimotor-association (S-A) axis expansion. A |** Distribution by timepoint and Yeo network of individual-level mean strength of functional connections, computed by averaging functional connectivity strength between each seed region and its 10% maximally functionally connected regions, further averaged across regions belonging to each of the seven Yeo networks; **B |** Group-level mean strength of functional connections (Fisher *r*-to-*z* values), computed by averaging regional individual-level mean strength of top 10% functional connections per seed region across all subjects; scatterplot displays patterns of mean strength of functional connections as a function of the seed region's loading on the adult S-A axis template (Margulies et al. 2016), tested by Spearman correlation and corrected for spatial autocorrelation using spin permutation testing (Alexander-Bloch et al. 2018), $r$ = -0.30, $p_{spin}$ = 0.010, color-coded by Yeo network; **C |** Estimated partial effects of age and pubertal stage on the mean strength of functional connections across Yeo networks; Both all effects (top panels) and only statistically significant effects (bottom panels) are displayed, at both the network- (left panels) and region-levels (right panels) (i.e., averaging connection strengths within each network or bin at the individual level); **D |** Estimated partial effects of mean strength of functional connections on S-A axis expansion; Both all effects and only statistically significant effects are displayed, at both the network- and region-levels (i.e., averaging connection strengths within each network or bin at the individual level). Boxplots are represented by boxes extending from the first to the third quartiles of the data with a line representing the median, whiskers extending from the boxes to the farthest data points within 1.5 times the interquartile range, and fliers representing data points past the ends of the whiskers. Statistical effects were computed with Big Additive Models (BAMs) including fixed effects of chronological age, pubertal stage, sex and total surface area, and random nested effects (subject, family, site). Lines represent the zero-centered component smooths extracted from the BAM models, illustrating the isolated contribution of each predictor to the response while accounting for all other model terms. Shaded ribbons denote 95% Bayesian credible intervals derived from the posterior distribution of the model coefficients, representing the localized uncertainty of the smooth estimate. V, visual; SM, somatomotor; DA, dorsal attention; VA, ventral attention; L, limbic; FP, frontoparietal; DMN, default-mode network. For analyses conducted at the region-level, regions were equally divided into 10 bins based on their position on the adult S-A axis (Margulies et al. 2016) and functional connectivity strength was averaged within each bin at the individual-level.

#### Sensitivity Analyses

To ensure our findings were robust to methodological choices and potential confounders, we conducted a series of sensitivity analyses.

We aimed to replicate our S-A axis expansion measures and related effects by recalculating S-A axis expansion at the regional level, using the sum of squared values of regional S-A axis loadings rather than network centroids, excluding outlier values that were 1.5 times greater than the third quartile and below the first quartile, based on this new metric ($N$ = 6122) (Supplementary Fig. 3A-C). We also assessed the impact of sample composition by re-analyzing the full dataset without excluding any S-A axis expansion outliers ($N$ = 6323) (Supplementary Fig. 3D-F). Our main results were broadly replicated across these analyses. One exception was the expansion-by-sex interaction on gradient flip probability, which was no longer statistically significant when using the regional expansion metric ($EDF$ = 1.15, $\chi^2$ = 1.56, $p$ = 0.098), suggesting that developmental trajectories are largely shared across sexes. Furthermore, in the full sample analysis, while significant effects persisted, trajectories exhibited more non-linearity and instability (Supplementary Fig. 3E-F). This variance was driven by a small subset of extreme, biologically improbable expansion values (as evidenced by deviation from main trajectories and widening 95% credibility intervals at high values of S-A axis expansion in Supplementary Fig. 3F), supporting our decision to exclude these outliers from the main analyses.

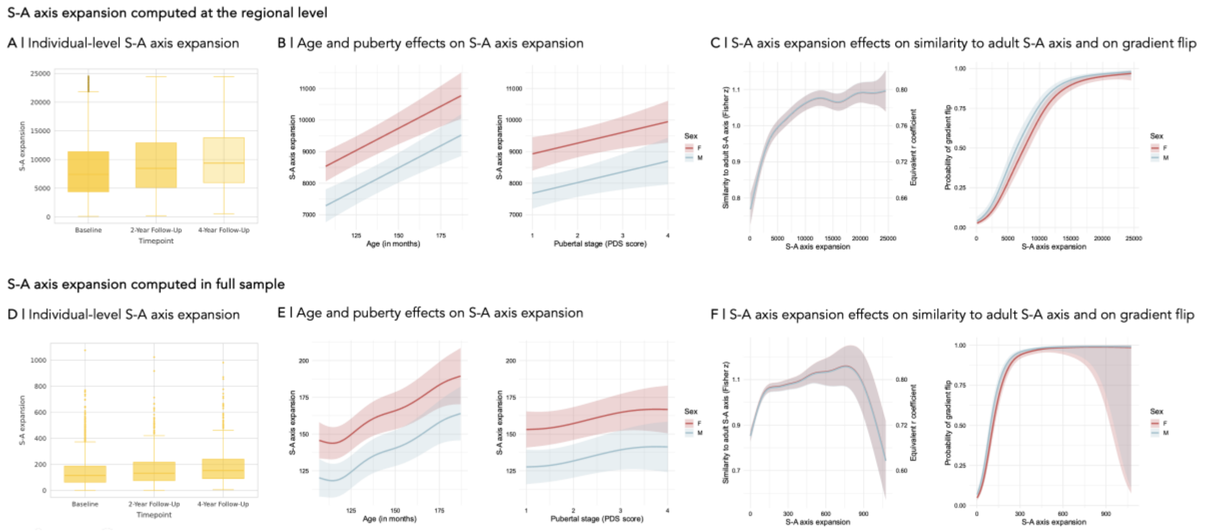


**Supplementary Figure 3. Sensorimotor-association (S-A) axis expansion and related effects, when computing the S-A axis at the regional level and in the full sample. A & D |** Distribution by timepoint of individual-level S-A axis expansion, defined as the sum of squared network centroids to capture total magnitude of diversion from S-A axis origin (0); **B & E |** Predicted marginal trajectories for age and pubertal stage effects on S-A axis expansion by sex; **C & F |** Predicted marginal trajectories for the S-A axis expansion effect on the similarity to adult S-A axis (model estimates are shown on the Fisher $r$-to-$z$ transformed scale; a secondary y-axis displays back-transformed Spearman $r$ values to ease interpretation) and the probably of gradient flip, by sex. Boxplots are represented by boxes extending from the first to the third quartiles of the data with a line representing the median, whiskers extending from the boxes to the farthest data points within 1.5 times the interquartile range, and fliers representing data points past the ends of the whiskers. Statistical effects were computed with Big Additive Models (BAMs) including fixed effects of chronological age, pubertal stage, sex and total surface area, and random nested effects (subject, family, site). Predictions were generated by holding all continuous covariates at their mean values and by excluding random nested effects to produce population-level trajectories. Shaded ribbons represent 95% Bayesian credible intervals derived from the posterior distribution of the model coefficients, representing the uncertainty in the population-level response by excluding the variance attributed to the random nested effects of subject, family, and site.

To ensure our developmental findings were not confounded by socioeconomic status (SES), we incorporated parental education as a covariate in our primary models, given its established impact on brain development (Tooley, Bassett, and Mackey 2021) (Supplementary Fig. 4). For this, we used the highest level of achieved education out of both parents if available, otherwise using the only reported level of achieved education. We excluded subjects with missing parental education data and our final sample included $N = 6319$ for full sample analyses, and $N = 6104$, for analyses excluding S-A axis expansion outliers.

Interestingly, while parental education was a significant predictor of gradient flip and functional connectivity profiles across nearly all networks (with the exception of the frontoparietal network), the inclusion of it as a covariate in our models did not notably alter our main results. Specifically, the significance and trajectories of all findings were largely replicated, with the sole exception being that the previously significant effect of pubertal stage on the length of functional connections in the default-mode network was attenuated to non-significance when controlling for parental education. This altogether suggests that our age- and pubertal status-dependent developmental findings are not driven by SES, even though SES is itself associated with some features of S-A axis development.

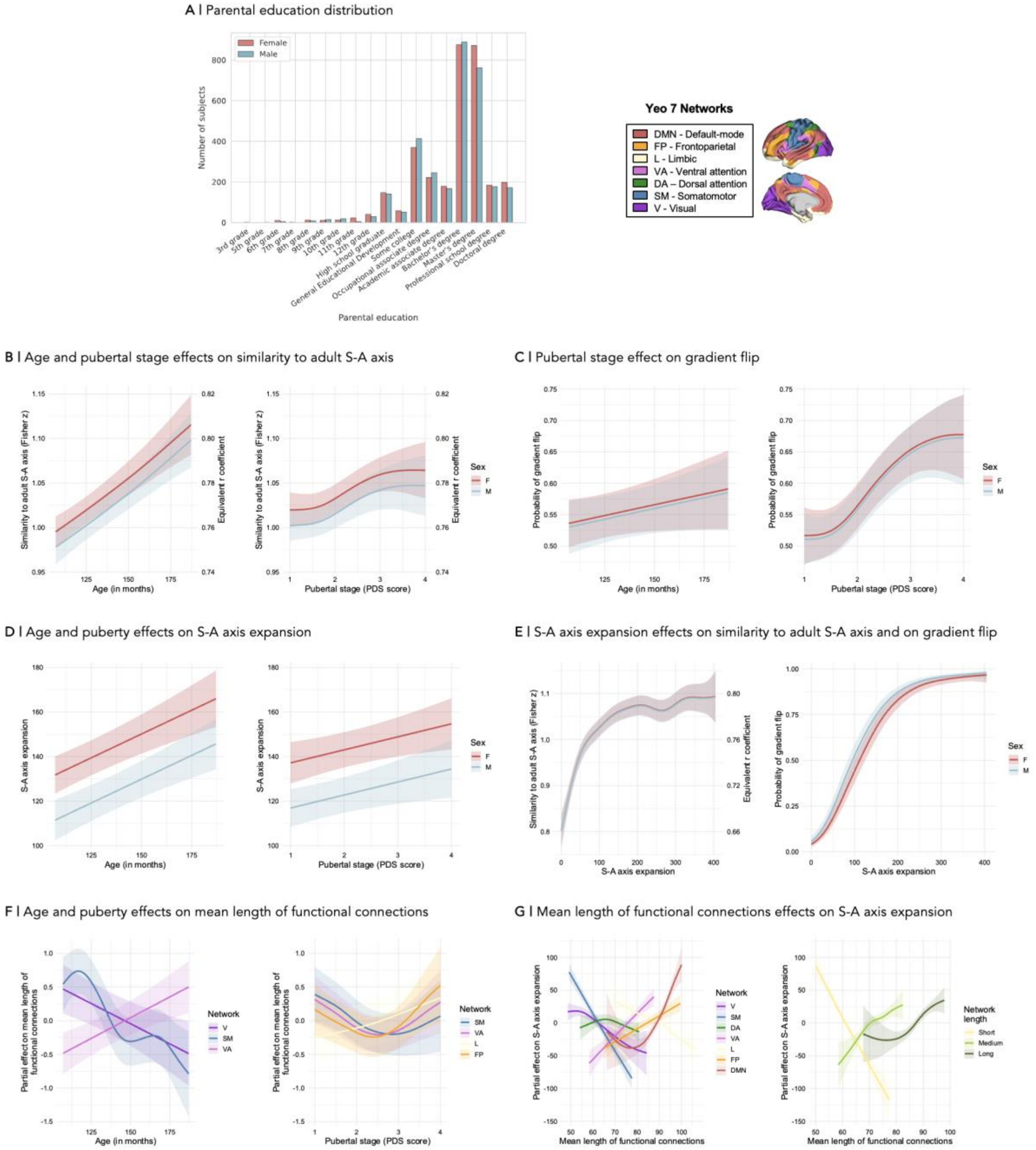


**Supplementary Figure 4. Statistical effects when controlling for a marker of socioeconomic status (SES), i.e., parental education. A |** Distribution of parental education by sex (of study participants; i.e., youth), representing the highest level of achieved education out of both parents if available, otherwise the only reported level of education; **B |** Predicted marginal trajectories for age and pubertal stage effects on the similarity to adult S-A axis by sex (model estimates are shown on the Fisher *r*-to-*z* transformed scale; a secondary y-axis displays back-transformed Spearman *r* values to ease interpretation); **C |** Predicted marginal trajectories for the age and pubertal stage effects on the probability of observing the gradient flip by sex (age effect is not statistically significant); **D |** Predicted marginal trajectories for age and pubertal stage effects on S-A axis expansion by sex; **E |** Predicted marginal trajectories for the S-A axis expansion effect on the similarity to adult S-A axis (model estimates are shown on the Fisher *r*-to-*z* transformed scale; a secondary y-axis displays back-transformed Spearman *r* values to ease interpretation) and the probably of gradient flip, by sex; **F |** Estimated partial effects of age and pubertal stage on the mean length of functional connections across Yeo networks; **G |** Estimated partial effects of mean length of functional connections on S-A axis expansion; Effects are estimated and visualized by individual Yeo network and by aggregating Yeo networks into categories based on their characteristic network length, i.e., short- (V, SM), mid- (DA, VA), and long- (L, FP, DMN) range networks. Statistical effects were computed with Big Additive Models (BAMs) including fixed effects of parental education, chronological age, pubertal stage, sex and total surface area, and random nested effects (subject, family, site). Predictions were generated by holding all continuous covariates at their mean values and by excluding random nested effects to produce population-level trajectories. Shaded ribbons represent 95% Bayesian credible intervals derived from the posterior distribution of the model coefficients, representing the uncertainty in the population-level response by excluding the variance attributed to the random nested effects of subject, family, and site.

To ensure our developmental findings were not confounded by body fat, given reports of associations between childhood obesity and pubertal timing (Aghaee et al. 2022), we incorporated body mass index (BMI) as a covariate in our primary models (Supplementary Fig. 5). BMI was conventionally calculated (kg/m2) and converted to a z-score according to age and sex (Gray, Schvey, and Tanofsky-Kraff 2020; Dehestani et al. 2024), based on the CDC 2000 Growth Charts and proposed metrics using the cdcanthro R package (https://github.com/CDC-DNPAO/CDCAnthro) (Freedman et al. 2020; Kuczmarski et al. 2002; Wei et al. 2020). We excluded subjects with missing anthropometric data or biologically implausible BMI z-scores following the recommendations of the SAS Program for CDC Growth Charts documentation (CDC 2024), namely z scores below -5 and above 10. Our final sample included $N = 6310$ for full sample analyses, and $N = 6095$ for analyses excluding S-A axis expansion outliers.

BMI was not a significant predictor of any of the tested developmental features, except functional connectivity profiles in the limbic network. Furthermore, its inclusion as a covariate in our models did not notably alter our main results, with the exception being that previously significant effect of pubertal stage on the length of functional connections in the limbic network was attenuated to non-significance, and the previously non-significant effect of pubertal stage on the length of functional connections in the visual network increased to reach significance. This altogether suggests that our age- and pubertal status-dependent developmental findings are not driven by BMI.

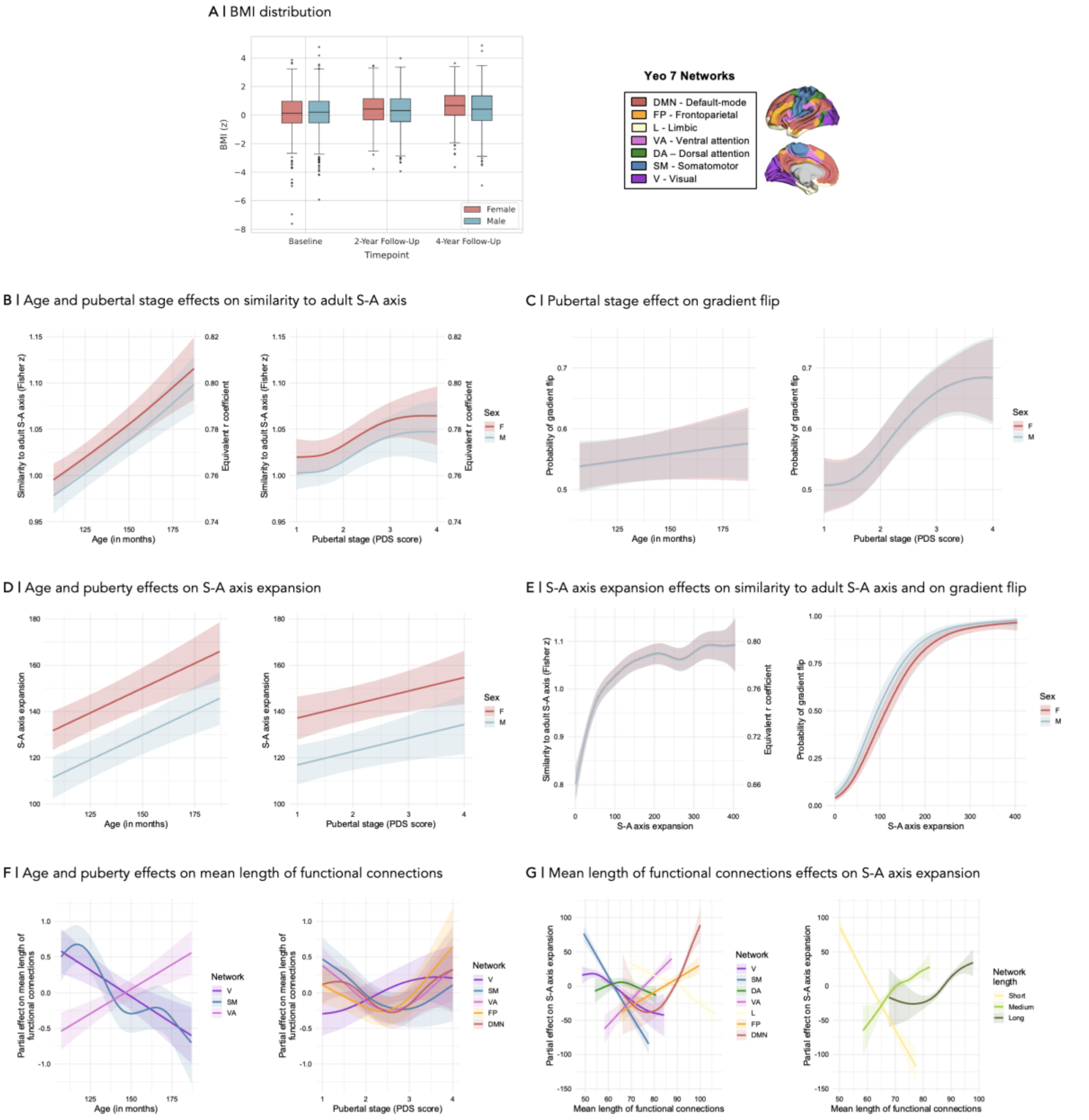


**Supplementary Figure 5. Statistical effects when controlling for a marker of body fat, i.e., body mass index (BMI). A |** Distribution of BMI values (z-scores according to age and sex, based on the CDC 2000 Growth Charts; (Kuczmarski et al. 2002) by sex and timepoint; **B |** Predicted marginal trajectories for age and pubertal stage effects on the similarity to adult S-A axis by sex (model estimates are shown on the Fisher *r*-to-*z* transformed scale; a secondary y-axis displays back-transformed Spearman *r* values to ease interpretation); **C |** Predicted marginal trajectories for the age and pubertal stage effects on the probability of observing the gradient flip by sex (age effect is not statistically significant); **D |** Predicted marginal trajectories for age and pubertal stage effects on S-A axis expansion by sex; **E |** Predicted marginal trajectories for the S-A axis expansion effect on the similarity to adult S-A axis (model estimates are shown on the Fisher *r*-to-*z* transformed scale; a secondary y-axis displays back-transformed Spearman *r* values to ease interpretation) and the probably of gradient flip, by sex; **F |** Estimated partial effects of age and pubertal stage on the mean length of functional connections across Yeo networks; **G |** Estimated partial effects of mean length of functional connections on S-A axis expansion; Effects are estimated and visualized by individual Yeo network and by aggregating Yeo networks into categories based on their characteristic network length, i.e., short- (V, SM), mid- (DA, VA), and long- (L, FP, DMN) range networks. Statistical effects were computed with Big Additive Models (BAMs) including fixed effects of BMIz, chronological age, pubertal stage, sex and total surface area, and random nested effects (subject, family, site). Predictions were generated by holding all continuous covariates at their mean values and by excluding random nested effects to produce population-level trajectories. Shaded ribbons represent 95% Bayesian credible intervals derived from the posterior distribution of the model coefficients, representing the uncertainty in the population-level response by excluding the variance attributed to the random nested effects of subject, family, and site.

We aimed to replicate our findings of functional connectivity profile effects (short-, medium-, and long-range network categories) on S-A axis expansion when computing the three categories of mean length of functional connections at the regional- rather than network-level. To this end, we assigned each region (rather than network) to one of the three categories (short, medium, long) based on the mean length of its top functional connections and then averaged across distances within each category. Our main results were broadly replicated across these analyses, with short-range regions still exhibiting a near-linear negative association with S-A axis expansion, and medium-to-long-range networks showing positive associations with S-A axis expansion. Here, effects appeared to be more linear and distinct than at the network level due to more homogeneous categorical definitions, given that they were based on actual mean length of regional functional connections as opposed to the characteristic lengths of connections made by functional networks (Supplementary Fig. 6). Detailed test statistics are provided in Supplementary Table 2.

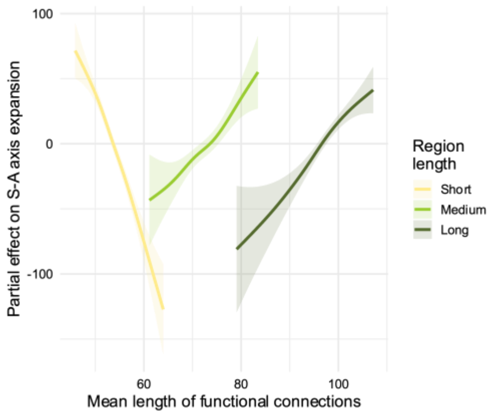


**Supplementary Figure 6. Mean length of functional connection effects on S-A axis expansion.** Estimated partial effects of mean length of functional connections on S-A axis expansion; Effects are estimated and visualized by aggregating Schaefer 400 regions into categories based on their mean length of connections at the individual level, i.e., by dividing regions into 3 quantiles and then averaging connection lengths within each quantile to characterize mean short-, mid-, and long- range regions. Statistical effects were computed with Big Additive Models (BAMs) including fixed effects of chronological age, pubertal stage, sex and total surface area, and random nested effects (subject, family, site). Predictions were generated by holding all continuous covariates at their mean values and by excluding random nested effects to produce population-level trajectories. Shaded ribbons represent 95% Bayesian credible intervals derived from the posterior distribution of the model coefficients, representing the uncertainty in the population-level response by excluding the variance attributed to the random nested effects of subject, family, and site.

#### Supplementary Methods

##### Participant inclusion

Participant inclusion was based on local availability at the time of analysis for ABCD Data Release 5.1.

MRI processing was restricted to individuals meeting the official ABCD Study quality control criteria for both T1-weighted (T1w) and resting-state fMRI (rs-fMRI) acquisitions. Specifically, we utilized the curated inclusion metrics provided by the ABCD Data Analysis, Informatics, and Control Center (DAIC) (Hagler et al. 2019), ensuring that only scans recommended for general use (as documented in the mri_y_qc_incl tabulated data) were included in our final sample.

We then applied our own inclusion criteria as shown in Supplementary Fig. 7. Namely, we retained subjects with a minimum of 2 runs of functional resting-state MRI scans (representing a minimum of 8 minutes of total scan time across runs) and with a complete mean functional connectivity matrix (i.e., no missing values). Then, we retained subjects with available parental Pubertal Development Scale (PDS) scores and total surface area.

Final lists of included subject IDs by timepoint are provided via GitHub to ensure full reproducibility.

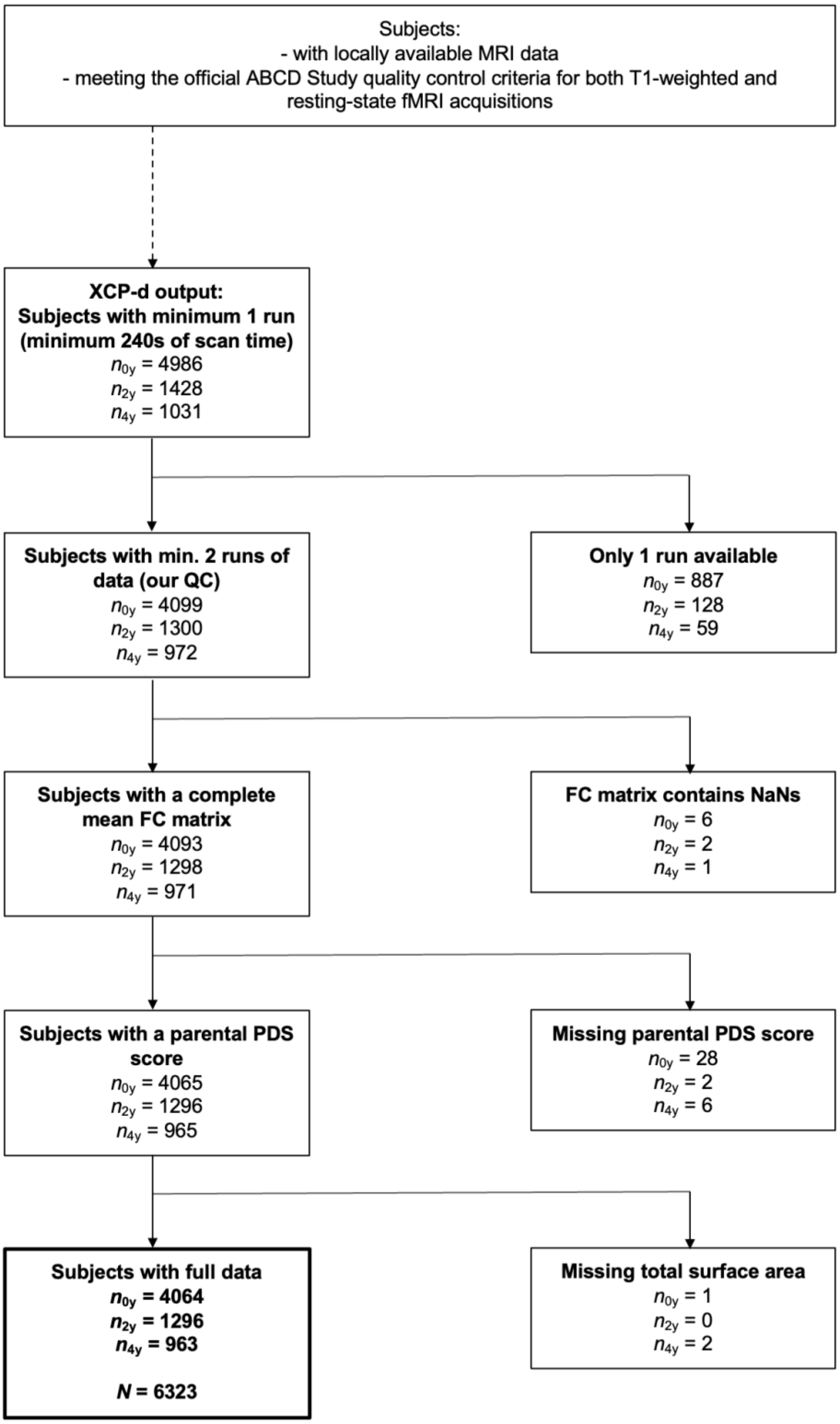


**Supplementary Figure 7. Participant Selection Flow Diagram.** Schematic representation of the data selection, outlining the exclusion criteria applied to the ABCD study sample and sample sizes by data collection timepoint: 0y, baseline; 2y, 2-year follow-up; 4y, 4-year follow-up. FC, functional connectivity; PDS, Pubertal Development Scale.

Hagler, Donald J., SeanN. Hatton, M. Daniela Cornejo, Carolina Makowski, Damien A. Fair, Anthony Steven Dick, Matthew T. Sutherland, B. J. Casey, Deanna M. Barch, Michael P. Harms, Richard Watts, James M. Bjork, Hugh P. Garavan, Laura Hilmer, Christopher J. Pung, Chelsea S. Sicat, Joshua Kuperman, Hauke Bartsch, Feng Xue, Mary M. Heitzeg, Angela R. Laird, Thanh T. Trinh, Raul Gonzalez, Susan F. Tapert, Michael C. Riedel, Lindsay M. Squeglia, Luke W. Hyde, Monica D. Rosenberg, Eric A. Earl, Katia D. Howlett, Fiona C. Baker, Mary Soules, Jazmin Diaz, Octavio Ruiz de Leon, Wesley K. Thompson, Michael C. Neale, Megan Herting, Elizabeth R. Sowell, Ruben P. Alvarez, Samuel W. Hawes, Mariana Sanchez, Jerzy Bodurka, Florence J. Breslin, Amanda Sheffield Morris, Martin P. Paulus, W. Kyle Simmons, Jonathan R. Polimeni, Andre van der Kouwe, Andrew S. Nencka, Kevin M. Gray, Carlo Pierpaoli, John A. Matochik, Antonio Noronha, Will M. Aklin, Kevin Conway, Meyer Glantz, Elizabeth Hoffman, Roger Little, Marsha Lopez, Vani Pariyadath, Susan RB. Weiss, Dana L. Wolff-Hughes, Rebecca DelCarmen-Wiggins, Sarah W. Feldstein Ewing, Oscar Miranda-Dominguez, Bonnie J. Nagel, Anders J. Perrone, Darrick T. Sturgeon, Aimee Goldstone, Adolf Pfefferbaum, Kilian M. Pohl, Devin Prouty, Kristina Uban, Susan Y. Bookheimer, Mirella Dapretto, Adriana Galvan, Kara Bagot, Jay Giedd, M. Alejandra Infante, Joanna Jacobus, Kevin Patrick, Paul D. Shilling, Rahul Desikan, Yi Li, Leo Sugrue, Marie T. Banich, Naomi Friedman, John K. Hewitt, Christian Hopfer, Joseph Sakai, Jody Tanabe, Linda B. Cottler, Sara Jo Nixon, Linda Chang, Christine Cloak, Thomas Ernst, Gloria Reeves, David N. Kennedy, Steve Heeringa, Scott Peltier, John Schulenberg, Chandra Sripada, Robert A. Zucker, William G. Iacono, Monica Luciana, Finnegan J. Calabro,

Duncan B. Clark, David A. Lewis, Beatriz Luna, Claudiu Schirda, Tufikameni Brima, John J. Foxe, Edward G. Freedman, Daniel W. Mruzek, Michael J. Mason, Rebekah Huber, Erin McGlade, Andrew Prescot, Perry F. Renshaw, Deborah A. Yurgelun-Todd, Nicholas A. Allgaier, Julie A. Dumas, Masha Ivanova, Alexandra Potter, Paul Florsheim, Christine Larson, Krista Lisdahl, Michael E. Charness, Bernard Fuemmeler, John M. Hettema, Hermine H. Maes, Joel Steinberg, Andrey P. Anokhin, Paul Glaser, Andrew C. Heath, Pamela A. Madden, Arielle Baskin-Sommers, R. Todd Constable, Steven J. Grant, Gayathri J. Dowling, Sandra A. Brown, Terry L. Jernigan, and Anders M. Dale. 2019. "Image Processing and Analysis Methods for the Adolescent Brain Cognitive Development Study." *NeuroImage* 202:116091. doi:10.1016/j.neuroimage.2019.116091.

Kuczmarski, Robert J., Cynthia L. Ogden, Shumei S. Guo, Laurence M. Grummer-Strawn, Katherine M. Flegal, Zuguo Mei, Rong Wei, Lester R. Curtin, Alex F. Roche, and Clifford L. Johnson. 2002. "2000 CDC Growth Charts for the United States: Methods and Development." *Vital and Health Statistics. Series 11, Data from the National Health Survey* (246):1–190.

Luo, Audrey C., Valerie J. Sydnor, Adam Pines, Bart Larsen, Aaron F. Alexander-Bloch, Matthew Cieslak, Sydney Covitz, Andrew A. Chen, Nathalia Bianchini Esper, and Eric Feczko. 2024. "Functional Connectivity Development along the Sensorimotor-Association Axis Enhances the Cortical Hierarchy." *Nature Communications* 15(1):3511.

Margulies, Daniel S., Satrajit S. Ghosh, Alexandros Goulas, Marcel Falkiewicz, Julia M. Huntenburg, Georg Langs, Gleb Bezgin, Simon B. Eickhoff, F. Xavier Castellanos, Michael Petrides, Elizabeth Jefferies, and Jonathan Smallwood. 2016. "Situating the Default-Mode Network along a Principal Gradient of Macroscale Cortical Organization." *Proceedings of the National Academy of Sciences* 113(44):12574–79. doi:10.1073/pnas.1608282113.

Serio, Bianca, Meike D. Hettwer, Lisa Wiersch, Giacomo Bignardi, Julia Sacher, Susanne Weis, Simon B. Eickhoff, and Sofie L. Valk. 2024. "Sex Differences in Functional Cortical Organization Reflect Differences in Network Topology Rather than Cortical Morphometry." *Nature Communications* 15(1):7714.

Sun, Lianglong, Tengda Zhao, Xinyuan Liang, Mingrui Xia, Qiongling Li, Xuhong Liao, Gaolang Gong, Qian Wang, Chenxuan Pang, Qian Yu, Yanchao Bi, Pindong Chen, Rui Chen, Yuan Chen, Taolin Chen, Jingliang Cheng, Yuqi Cheng, Zaixu Cui, Zhengjia Dai, Yao Deng, Yuyin Ding, Qi Dong, Dingna Duan, Jia-Hong Gao, Qiyong Gong, Ying Han, Zaizhu Han, Chu-Chung Huang, Ruiwang Huang, Ran Huo, Lingjiang Li, Ching-Po Lin, Qixiang Lin, Bangshan Liu, Chao Liu, Ningyu Liu, Ying Liu, Yong Liu, Jing Lu, Leilei Ma, Weiwei Men, Shaozheng Qin, Jiang Qiu, Shijun Qiu, Tianmei Si, Shuping Tan, Yanqing Tang, Sha Tao, Dawei Wang, Fei Wang, Jiali Wang, Pan Wang, Xiaoqin Wang, Yanpei Wang, Dongtao Wei, Yankun Wu, Peng Xie, Xiufeng Xu, Yuehua Xu, Zhilei Xu, Liyuan Yang, Huishu Yuan, Zilong Zeng, Haibo Zhang, Xi Zhang, Gai Zhao, Yanting Zheng, Suyu Zhong, and Yong He. 2025. "Human Lifespan Changes in the Brain's Functional Connectome." *Nature Neuroscience* 28(4):891–901. doi:10.1038/s41593-025-01907-4.

Taylor, Hoyt Patrick, Khoi Minh Huynh, Kim-Han Thung, Guoye Lin, Wenjiao Lyu, Weili Lin, Sahar Ahmad, and Pew-Thian Yap. 2026. "Functional Hierarchy of the Human Neocortex across the Lifespan." *Nature* 1–10. doi:10.1038/s41586-026-10219-x.

Tooley, Ursula A., Danielle S. Bassett, and Allyson P. Mackey. 2021. "Environmental Influences on the Pace of Brain Development." *Nature Reviews Neuroscience* 22(6):372–84. doi:10.1038/s41583-021-00457-5.

Wei, Rong, Cynthia L. Ogden, Van L. Parsons, David S. Freedman, and Craig M. Hales. 2020. "A Method for Calculating BMI Z-Scores and Percentiles above the 95th Percentile of the CDC Growth Charts." *Annals of Human Biology* 47(6):514–21. doi:10.1080/03014460.2020.1808065.

Yeo, BT Thomas, Fenna M. Krienen, Jorge Sepulcre, Mert R. Sabuncu, Danial Lashkari, Marisa Hollinshead, Joshua L. Roffman, Jordan W. Smoller, Lilla Zöllei, and Jonathan R. Polimeni. 2011. "The Organization of the Human Cerebral Cortex Estimated by Intrinsic Functional Connectivity." *Journal of Neurophysiology*.
